# An engineered closed-shell, two-component, 480-subunit nucleocapsid

**DOI:** 10.1101/2025.10.22.683813

**Authors:** Mikail D. Levasseur, Naohiro Terasaka, Angela Steinauer, Stephan Tetter, Sara Pfister, Beat H. Meier, Donald Hilvert

**Author notes:** Corresponding Author: **Donald Hilvert** − *Laboratory of Organic Chemistry, ETH Zurich, 8093 Zurich, Switzerland. M.D.L. and N.T. contributed equally. Earth-Life Science Institute, Institute of Future Science, Institute of Science Tokyo, 2-12-1 Ookayama, Meguro-ku, Tokyo, 152-8550 Japan (Naohiro Terasaka). Laboratory of Biomolecular Engineering and Nanomedicine, EPFL Lausanne, 1015 Lausanne, Switzerland. MRC Laboratory of Molecular Biology, Cambridge, UK.

## Abstract

Self-assembling protein cages are powerful nanoscale containers for biotechnology and medical applications. Two-component systems are especially attractive due to their potential for functional complexity. In this study, we demonstrate that the subunits of the 240mer nucleocapsid NC-4, which was previously evolved to package and protect its encoding mRNA, can be split into two fragments without disrupting cage assembly or structure, generating a two-component, 480-subunit capsid. This modification introduces additional termini on the cage’s exterior surface, creating new opportunities for functionalization. We exploited these new sites by genetically appending peptide and protein tags to the exterior surface of split NC-4 (spNC-4), enabling site-specific glycosylation via post-translational modification and cell-specific delivery by targeted antibody recruitment. Our findings broaden the utility of the NC-4 nucleocapsid. By extension, splitting related protein compartments that bind diverse cargoes could offer a robust platform for biotechnological applications requiring simultaneous encapsulation and customizable surface modification.

**SIGNIFICANCE STATEMENT:** Natural protein shells such as viral capsids and bacterial microcompartments have inspired efforts to design synthetic compartments that protect and deliver functional molecules. Here we show that a nonviral, artificially evolved nucleocapsid can be split into two fragments that reassemble into a closed, two-component, 480-subunit cage that packages its own mRNA. This redesign preserves the original architecture and selective RNA packaging while creating new engineerable sites on both the interior and exterior surfaces, enabling simultaneous control of internal cargo binding and external modification. The resulting two-component architecture highlights the structural plasticity of synthetic nucleocapsids and provides a general strategy for constructing modular, evolvable protein containers for biotechnology and synthetic biology.

## INTRODUCTION

Closed-shell protein assemblies—such as viral capsids, ferritins, and bacterial microcompartments (BMCs)—are remarkable natural structures that serve a wide range of biological functions, including packaging, protecting, and transporting nucleic acids,^1^ templating the formation of nanoparticles,^2^ and facilitating compartmentalized metabolic cascades.^3^ The inherent engineerability of such protein shells has enabled their repurposing for applications beyond their native roles, for example, in the development of improved vaccines, targeted delivery vehicles, and versatile scaffolds for synthetic biology.^4–7^

Inspired by such systems, novel protein assemblies have been created either through computational de novo design^8–11^ or the redesign and directed evolution of natural systems.^12^ A recent example of the latter approach is NC-4, a nonviral nucleocapsid we designed to package its encoding mRNA.^13^ NC-4 originated from the bacterial shell-forming enzyme lumazine synthase, which we modified by circular permutation to relocate its N- and C-termini to the lumen and addition of an N-terminal RNA-binding peptide called λN+ (Figure S1).^14^ The construct was then subjected to multiple rounds of directed evolution involving mutagenesis and increasingly stringent selection. Over successive generations, these selective pressures led to the emergence of NC-4, which features a highly regular, interlaced 240-subunit *T*=4 icosahedral architecture, enhanced nuclease resistance, and a robust RNA packaging cassette in its mRNA, resulting in efficient and specific encapsidation of its own mRNA over host nucleic acids.

To further expand NC-4’s utility, genetic modification of its exterior surface would be desirable. One common approach for modifying virus-like particles involves insertion of heterologous sequences into surface-exposed loops,^15–19^ which works well for short peptides but is generally unsuitable for larger proteins unless their termini are close together. Moreover, inserting large or complex sequences into loops can destabilize the subunits or interfere with their proper assembly and function. In contrast, splitting the NC-4 subunits by cutting through an exterior loop would generate two fragments, whose termini could be used for further functionalization. This strategy, which has been successfully applied to the hepatitis B virus capsid protein^20^ and the artificial fusion-protein nanocage TIP60,^21^ offers several advantages: each fragment can be engineered separately, allowing for independent modification of interior and exterior surfaces; it enables the incorporation of larger or more structurally complex domains that would not fit within a loop; and it allows for conditional assembly and functional control, as the fragments can be designed to reassemble only under specific conditions. As a result, splitting provides a potentially more versatile and modular approach for engineering complex, multifunctional protein assemblies.

In the current study, we show that by introducing a break in the exterior NC-4 loop used for circular permutation in the original nucleocapsid design, the NC-4 subunits can be split into two fragments that still spontaneously assemble in the cytosol of *Escherichia coli* to form a two-component, 480-subunit *T*=4 capsid. This split system not only adopts a structure closely related to the original NC-4 capsid but also retains the ability to selectively bind its encoding mRNA, even within the complex, RNA-rich environment of the cytosol. Unlike the original NC-4, where both termini are internal, each subunit of the split protein now has one terminus on the inside and one on the outside. This arrangement enables independent modification of both the interior and exterior surfaces of the capsid, for example, attaching peptide tags like λN+ on the interior for cargo binding, while using different tags on the exterior for chemical modification, targeting, or other applications.

## RESULTS

### Design of split nucleocapsid spNC-4

To create a split version of NC-4 (spNC-4), we designed a bicistronic construct encoding an N subunit (residues Met1 to Ser113) and a C subunit (residues Ile114 to Gly197) for co-expression in *E. coli* (Figure 1A). The N subunit includes both the λN+ peptide and a hexahistidine sequence at its N- and C-termini, respectively, whereas the C subunit lacks an affinity tag. A T7 promoter drives transcription of the genes encoding both fragments, and each gene is preceded by a ribosome binding site required for translation. The boxB packaging signals^22^ that had been introduced in the 5′ and 3′ untranslated regions (UTRs) of the NC-4 gene were also retained to promote mRNA encapsulation. To enable translation of the newly created C subunit, six nucleotides (5′-ATGGGA), including the ATG start codon, were inserted at the 5′ end of the corresponding gene. As a consequence, the C subunit possesses two additional amino acids, an initiator Met followed by a Gly, that were not present in the original NC-4 sequence. To keep the sequence numbering consistent with that of NC-4, we refer to the inserted amino acids in spNC-4 as Met113A and Gly113B.

**Figure 1.**
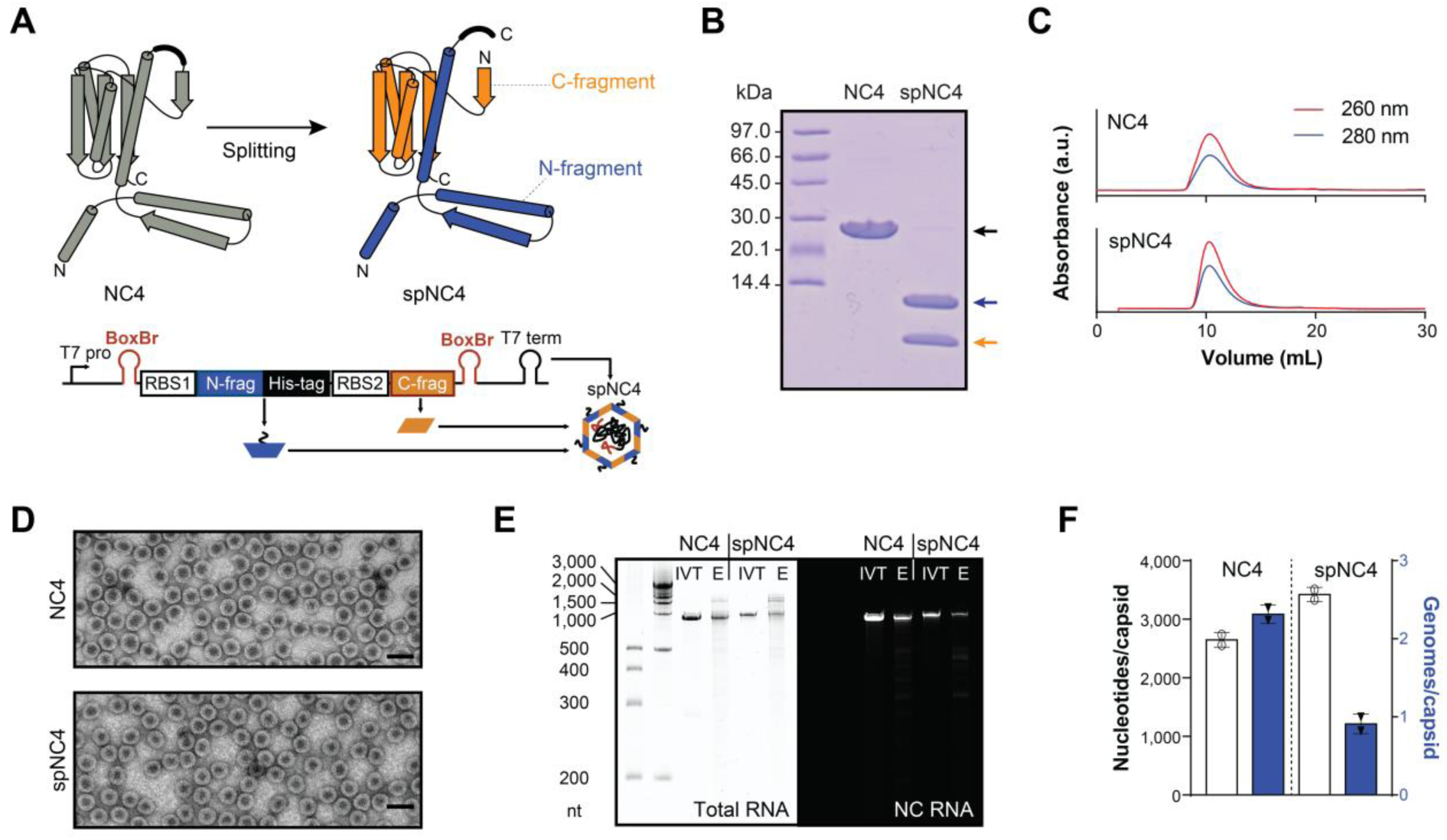
Design and characterization of spNC-4. (**A**) Structural models illustrating the splitting of an NC-4 monomer (top) and the corresponding topology diagram of the gene encoding the two spNC-4 fragments (bottom). Both components coassemble into cages around their mRNA via packaging signals such as the BoxBr tag. (**B**) Coomassie-stained SDS-PAGE of NC-4 and spNC-4. The NC-4 band is indicated by a black arrow, and the N- and C-fragments of spNC-4 are indicated by blue and orange arrows, respectively. (**C**) Size-exclusion chromatography of NC-4 and spNC-4 cage variants. Analytical profiles of the purified assemblies are shown, with absorbance at 260 nm (nucleic acid) and 280 nm (protein) plotted as solid red and blue lines, respectively. (**D**) Negative stain transmission electron micrographs of NC-4 (top) and spNC-4 (bottom). Scale bars, 50 nm. (**E**) 5% PAGE of NC-4 and spNC-4 stained either for total RNA with GelRed (left) or with the fluorogenic dye DFHBI-1T, which selectively binds the broccoli aptamer “BoxBr” (right). IVT, in vitro-transcribed reference mRNA; E, extracted RNA. (**F**) Nucleotide content per capsid (white bars) was determined from 260/280-nm absorbance ratios. The fraction of total extracted RNA corresponding to cage-encoding mRNA (blue bars) was quantified by RT-qPCR. Data are presented as means ± SD from two biological replicates.

### Biochemical properties of spNC-4

The two spNC-4 subunits were co-expressed in *E. coli* from the bicistronic gene described above. Assembled particles were isolated from the cells and purified by metal-ion affinity chromatography, followed by size exclusion chromatography (SEC) and anion exchange chromatography. Analysis of the purified nucleocapsids by denaturing polyacrylamide gel electrophoresis (PAGE) verified that the purified particles contain two protein chains of ∼12 and ∼9 kDa. For comparison, only a single protein of ∼21 kDa is observed for the parent NC-4 capsid (Figure 1B). Reinjection of the purified capsids on an analytical SEC column revealed that spNC-4 and NC-4 have similar sizes, eluting near the exclusion volume of the column (Figure 1C). The yields of the purified nucleocapsids per liter culture were also comparable for NC-4 (∼35 mg) and spNC-4 (∼38 mg). In negatively stained transmission electron micrographs, the two capsids were indistinguishable, with spherical shells of 32 ± 1 nm diameter (Figure 1D). These data show that cleavage of the external loop in NC-4 to create the two-component spNC-4 cage does not adversely affect capsid yield, size, or morphology.

The parent nucleocapsid NC-4 packages two to three copies of its encoding mRNA when assembled in *E. coli*. To examine how protein chain disconnection affected RNA packaging efficiency, we extracted RNA from both NC-4 and spNC-4 and analyzed it on a denaturing urea PAGE gel. Staining total RNA with GelRed showed that NC-4 exhibited one prominent band that coincided with the in vitro-transcribed 863-nt control mRNA (Figure 1E, left gel). The faint bands running at higher molecular mass between 1,500 and 3,000 nucleotides were previously assigned to ribosomal RNA (rRNA).^13^ For spNC-4, a slight shift to higher molecular mass was observed for the reference RNA, reflecting the presence of the 76 nt-long insert between the open reading frames encoding the two protein subunits. In addition, the fraction of host rRNA to capsid mRNA increased relative to NC-4 (60.3% to 12.0 %, calculated from band intensities in Figure 1E, left gel). Staining the same gel with the fluorogenic dye DFHBI-1T, which binds Broccoli aptamers introduced with the BoxB tags,^23^ confirmed that spNC-4 encapsidates less target mRNA than NC-4 (Figure 1E, right gel). Real-time quantitative polymerase chain reaction (RT-qPCR) was used to quantify these qualitative differences. The total number of nucleotides per particle was first established by UV absorbance, and the fraction of the extracted RNA corresponding to the target transcript was then determined by qPCR (Figure 1F). This analysis revealed that spNC-4 encapsulates 29.1% more total RNA than NC-4 (3421 ± 124 nt to 2648 ± 124 nt), but the number of target mRNAs decreases from 2.3 ± 0.1 to 0.9 ± 0.1 per capsid.

### Structural characterization of spNC-4

During its design and evolution, NC-4 underwent major structural changes. While parental lumazine synthase naturally assembles as a *T*=1 dodecahedron composed of 12 pentamers,^24^ NC-4 instead adopts a novel *T*=4 architecture.^13^ The altered assembly state arose from a domain swap that generated new trimeric building blocks. The resulting structure not only increased the particle size (240 versus 60 subunits) but also enhanced nuclease resistance, owing to its smaller pore size compared to other lumazine synthase derivatives. Because lumazine synthase scaffolds are known to undergo structural rearrangements in response to mutation,^12^ it was critical to assess whether the split-cage variant would preserve the NC-4 architecture.

We analyzed spNC-4 samples by cryogenic electron microscopy (cryoEM) and single-particle reconstruction. Over 2,000 movies were collected on a 300 kV microscope. An average of ∼72,000 particles was classified in 2D, and good, symmetric classes (∼63,000 particles) were further classified in 3D. Initial three-dimensional models with icosahedral symmetry were built de novo from the data to avoid bias from the NC-4 structure. The final map, refined to 3.1 Å (Table S1), is superimposable on that of the capsid formed by the non-split protein (Figure 2).

**Figure 2.**
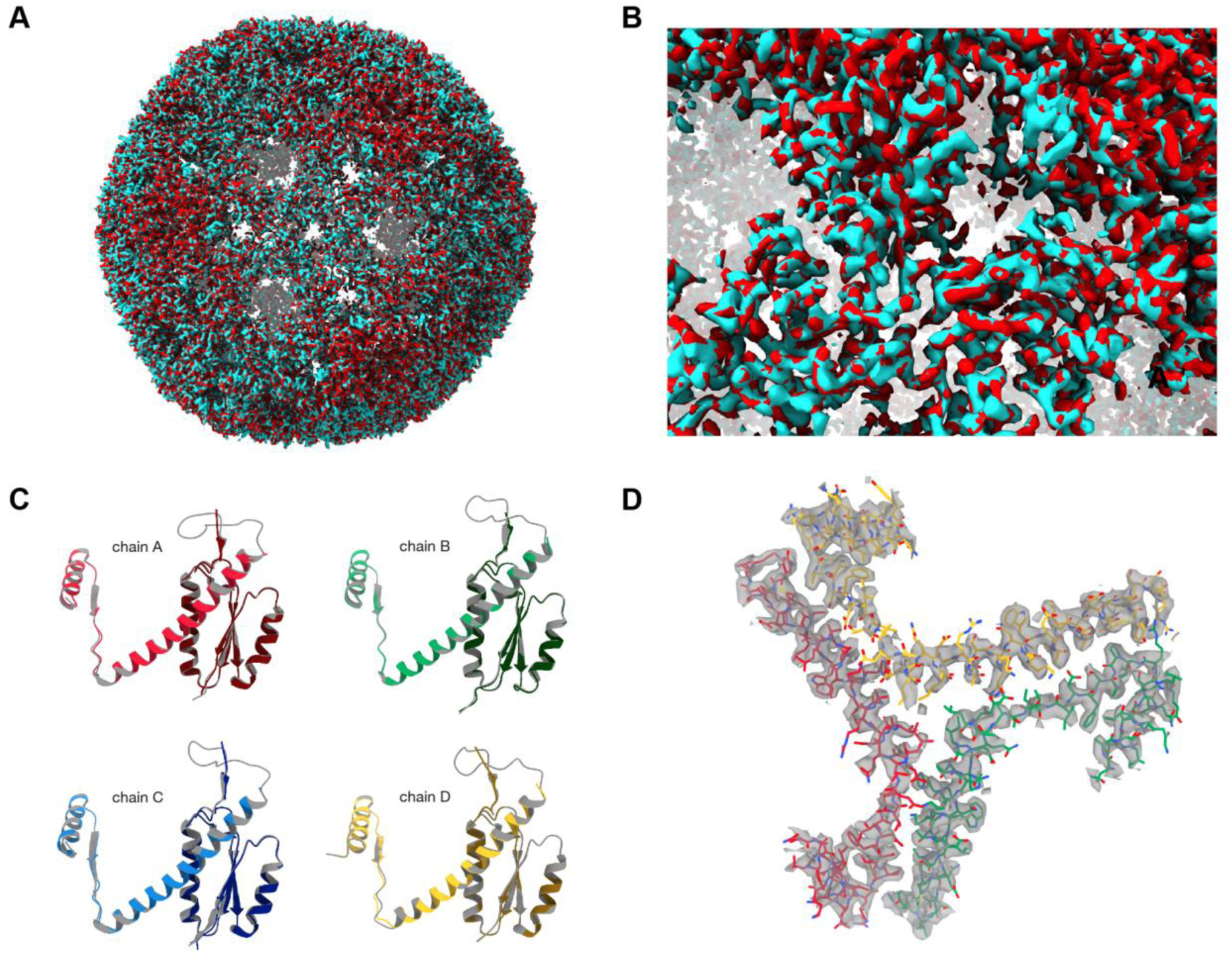
NC-4 maintains its structure after splitting. (**A**) Structural overlay of spNC-4 (cyan) and NC-4 (EMD-11635, red). (**B**) Close-up view of the overlay highlighting the overall structural correspondence. (**C**) Cartoon representations of spNC-4 chains superimposed on the corresponding quasi-equivalent subunits of NC-4 demonstrate that backbone conformation is preserved upon splitting. (**D**) Three quasi-equivalent N fragments are shown with the model fitted into the experimentally determined density (gray). The intersubunit domain swap of the parent NC-4 capsid is retained, with each N-fragment complementing two different C-fragments.

The atomic model constructed using the NC-4 template and refined against the spNC-4 map demonstrates that splitting did not alter the overall protein fold. Backbone topology and the majority of sidechain orientations are maintained in the split construct. Even the hinge element that enabled the domain swap is preserved in the N subunit after splitting. This is particularly notable as the original domain swap partitioned the wild-type lumazine synthase fold between two polypeptide chains, and the current split further divides the globular core into three separate polypeptides. The only clear difference between the structures is localized—as expected—in the external NC-4 loop (Thr103-Ile116) that was cleaved to create spNC-4. This region is poorly resolved in NC-4, indicating a flexible and poorly structured element, but becomes undetectable in spNC-4, suggesting substantially increased floppiness.

Solid-state nuclear magnetic resonance (NMR), which has provided valuable insights into the structure and dynamics of viral capsids,^25, 26^ was also used to probe potential differences between the spNC-4 and NC-4 protein cages. For these experiments, spNC-4 and NC-4 were uniformly labeled with ^15^N and ^13^C isotopes by production in *E. coli*.^27^ The purified samples were sedimented into solid-state NMR rotors by ultracentrifugation,^28, 29^ and solid-state NMR magic-angle spinning (MAS) experiments were carried out as described previously for hepatitis B virus.^30–32^

For the assignment of ^1^H, ^13^C, and ^15^N chemical shifts, a series of 2-dimensional and 3-dimensional spectra were recorded as summarized in Table S2. The spNC-4 construct was analyzed first, and the assignments were subsequently transferred to the parent NC-4 nucleocapsid. We successfully assigned 61% of the ^1^H, ^13^C, and ^15^N resonances for spNC-4 (backbone, minus the flexible N-terminus), and 66% for NC-4 (Tables S3-S5). Some residues could not be assigned for either protein, in most cases because the corresponding peaks were absent, especially at the N-terminus and in the outer loop region. The flexibility of these regions of the protein likely results in broad NMR peaks with intensities below the detection limit under the experimental conditions.^32^

Almost all peaks in the spNC-4 spectra are directly superimposable on their counterparts in the NC-4 spectra (Figure 3), as expected given the similarities of their respective cryo-EM structures. Nonetheless, the spectra differ in two respects. First, peaks for residues Ile114, Glu115, and, to some extent, Ile116 are observed for spNC-4, but not for NC-4. In spNC-4, these residues are proximal to the cut site, immediately following Met113A and Gly113B at the N-terminus of the C subunit. In contrast, in NC-4, they are part of the exterior loop that was introduced for circular permutation, which adopts four different conformations in the four quasi-equivalent subunits in the *T*=4 icosahedral lattice. Conceivably, the structural heterogeneity in this poorly resolved region caused peak splitting in the solid-state NMR spectra of NC-4, which, in combination with the dynamics of the loop, resulted in a signal that was below the detection limit.

**Figure 3.**
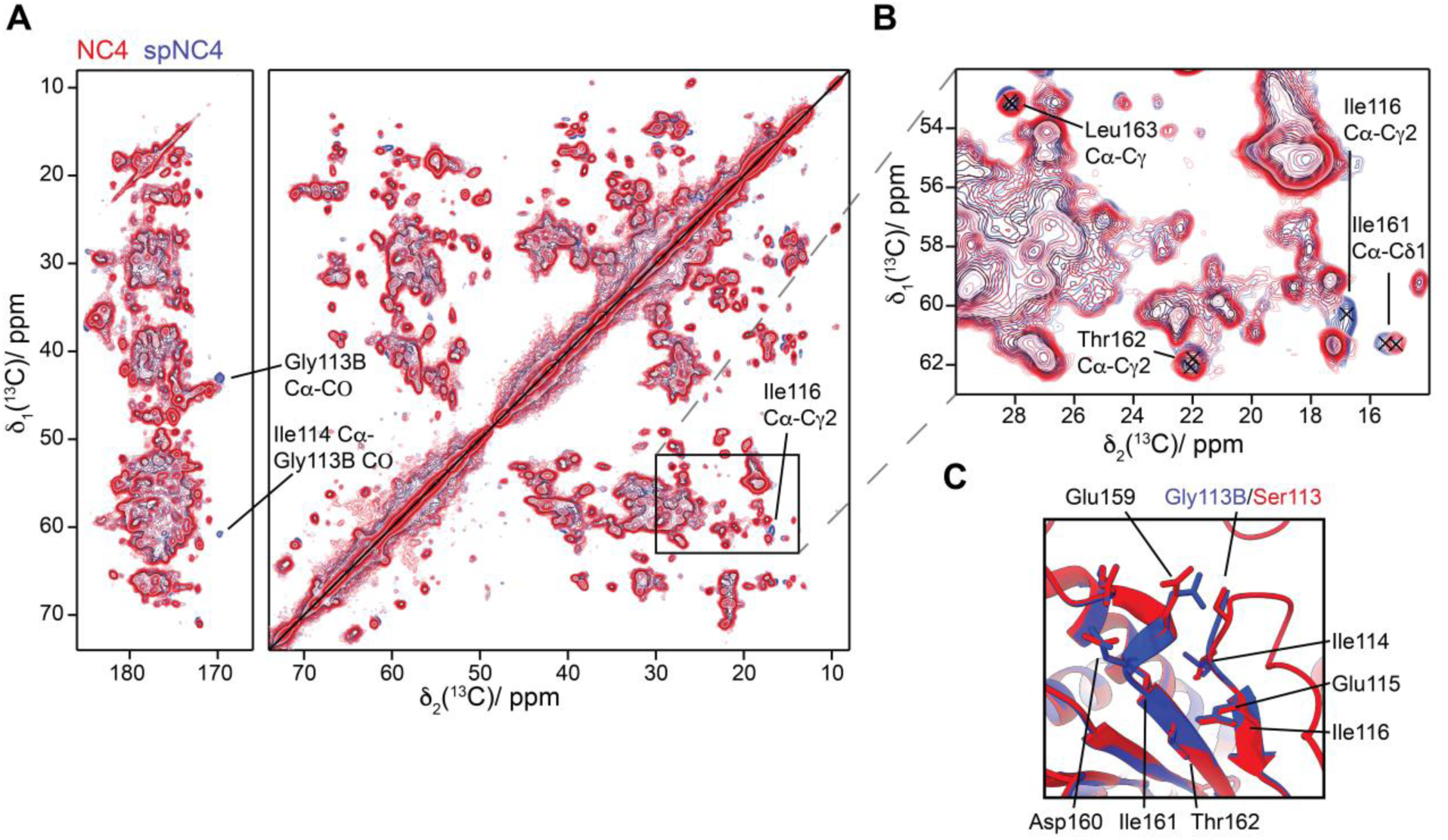
Superposition of the ^13^C-^13^C DARR spectra for NC-4 (red) and spNC-4 (blue). Spectra were acquired at 850 MHz, 17 kHz MAS, using a 3.2 mm rotor for a 20 ms DARR mixing time. Residue-specific signals are colored according to their parent structure: NC-4 residues are in red, spNC-4 in blue. Major spectral differences in panels (**A**) and (**B**) are labeled with the corresponding atom. Residues Ser113–Ile116 are highlighted in panel **C** as these are visible in spNC-4 but not in NC-4 spectra, as shown in panels A and B; residues Glu159-Thr162, which are proximal to the cleavage site and showed small chemical shift changes, are also illustrated. Additional examples of chemical shift changes are provided in Figures S3–S7. For clarity, see Figure S8 for an overlay with the blue spectrum displayed on top of the red spectrum.

The second difference concerns chemical-shift changes. The chemical shifts of nuclei are highly sensitive to their local electronic environment and can report on subtle changes in protein dihedral angles.^33^ The differences in chemical shifts between spNC-4 and NC-4 are generally small (<0.3 ppm), often within the range of estimated standard deviations. Though small in relative terms, the Cα and Cβ atoms of residues Glu159–Ile161 show the biggest changes (Figure S2), with Asp160 exhibiting the largest absolute differences. The sidechains of Asp160 and Arg157 are located on the exterior surface of the capsid, oriented towards each other to form a salt bridge. The side chain orientations of these residues are not well-resolved in the spNC-4 cryo-EM structure, however, and the chemical-shift change suggests that this electrostatic interaction may be perturbed in the split capsids. This hypothesis is bolstered by the analysis of chemical shift differences for C’, N, amide H, and sidechain carbons between the two protein cages (Figures S3-S7), which are largest for residue Glu159 in the same protein segment.

### Glycosylation of spNC-4 via external peptide glycosylation motifs

A common strategy for functionalizing protein cages involves appending short peptide tags^34, 35^ or entire folded proteins^36^ to the exposed N-or C-terminus on the exterior surface of the particles. To test whether spNC-4 can be similarly modified without disrupting its assembly, we genetically added a glycosylation tag to the N subunit of spNC-4, immediately preceding the C-terminal hexahistidine tag (Figure 4A). We call the resulting construct spNC-4-GI. The 17 amino acid-long tag (GSGSGAHATA<u>NAT</u>AHAS) consists of a flexible linker followed by a recognition sequence for *Actinobacillus pleuropneumoniae* asparagine N-glucosyltransferase (ApNGT), which catalyzes the transfer of a single β-linked glucose to the asparagine residue in the consensus sequon (underlined).^37, 38^

**Figure 4.**
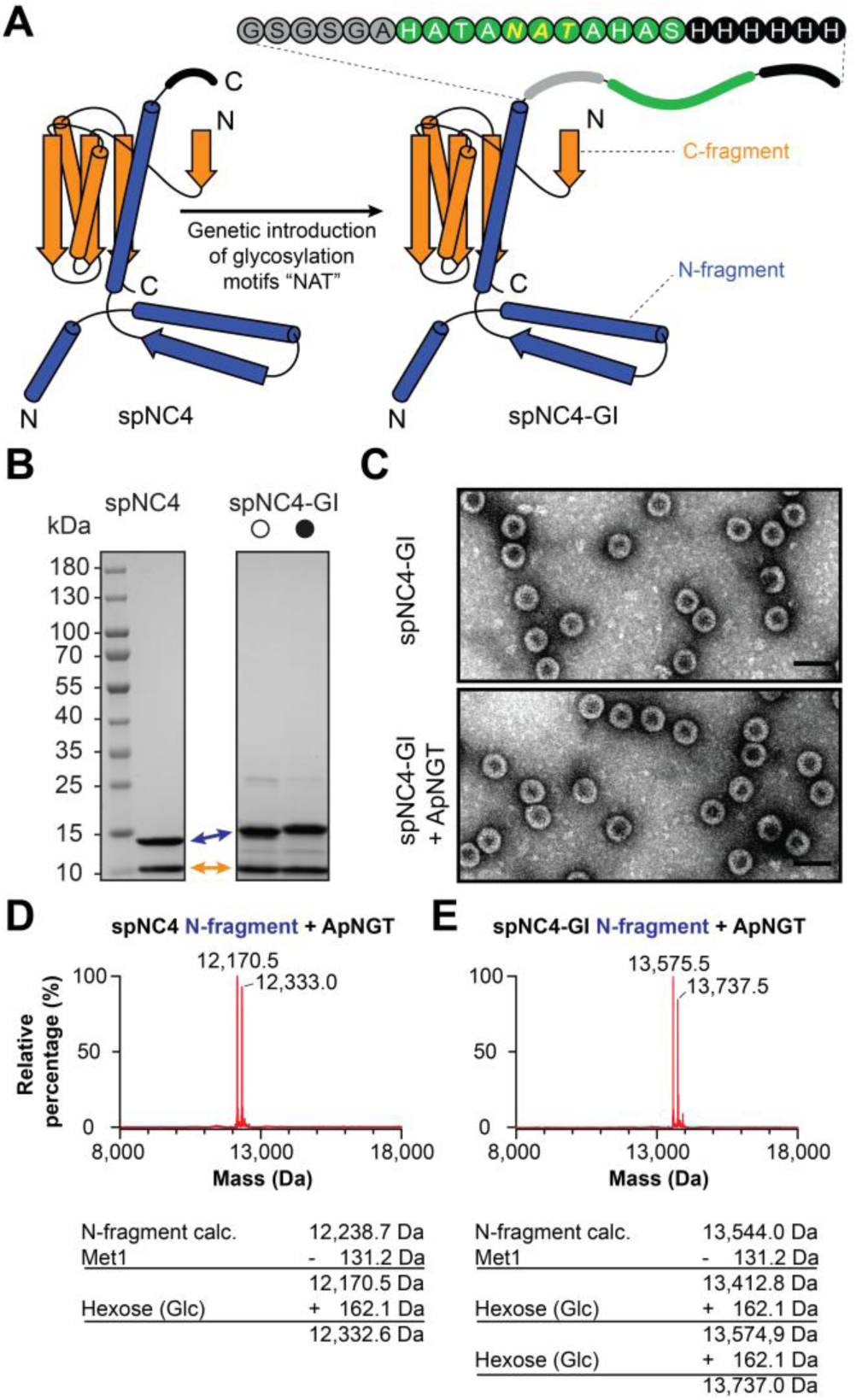
Glycosylation of spNC-4. (**A**) Structural model of an spNC-4 monomer showing the engineered insertion of the glycosylation motif “NAT” (yellow residues) at the outward-facing C-terminus, generating the variant spNC-4-GI. The glycosylation sequon was introduced within a flexible tag (green spheres) positioned between an amino acid spacer (gray spheres) and a hexahistidine tag (black spheres). (**B**) Coomassie-stained SDS-PAGE analysis of spNC-4-GI expressed in the absence (open circles) or presence (filled circles) of the glucosyltransferase ApNGT. The spNC-4 N-and C-fragments are indicated by blue and orange arrows, respectively. Upon co-expression with ApNGT, a mobility shift of the tagged N-fragment band is observed, consistent with glycosylation.(**C**) TEM analysis of spNC-4-GI without (top) and with (bottom) ApNGT co-expression. (**D**) MS analysis of the N-fragment of spNC-4 (left) and spNC-4-GI (right) following co-expression with ApNGT.

Glycosylation of spNC-4-GI capsids was investigated by co-producing the split protein with ApNGT in *E. coli*. Intact cages were isolated and purified by affinity chromatography and analyzed by SDS-PAGE, transmission electron microscopy (TEM), and mass spectrometry (MS) (Figure 4B-D). The resulting particles were similar in size to spNC-4-GI produced in the absence of ApNGT and the parent NC-4 nucleocapsid (spNC-4-GI + ApNGT diameter = 32.4 ± 1.1 nm; spNC-4-GI diameter = 31.3 ± 0.8 nm; NC-4 diameter = 32.0 ± 1.0 nm) (Figure 4C). Mass spectrometric analysis showed, however, that the N subunit of spNC-4-GI was quantitatively glycosylated, affording a 60:40 mixture of singly and doubly modified subunits (Figure 4E). For samples produced under the same conditions but in the absence of the glucosyltransferase, no glycosylation was observed (Figures 4B and S9). Since the glycosylation tag only contains a single asparagine, these results suggest that the N subunit must possess a second glycosylation sequon. Consistent with this hypothesis, when spNC-4 lacking the glycosylation tag was coproduced with ApNGT, ∼50% of its N subunits were also modified with a single hexose unit (Figure 4D).

Proteomics analysis of the glycosylated spNC-4-GI sample (Tables 1 and 2) confirmed essentially quantitative (99.5%) addition of a hexose unit to the NAT sequon in the tag on the N subunit. As a result, each fully assembled nucleocapsid presents 240 copies of the monosaccharide on its exterior surface of the assembled spNC-4-GI nucleocapsids. In addition to the targeted glycosylation site, two surface-exposed residues that project into the interior of the capsid—Asn39 (53%) and, to a lesser extent, Asn49 (7%)—were also modified in both spNC-4-GI and spNC-4. Asn39 is located in a consensus sequon (NAS) for ApNGT, whereas Asn49 is found in a less preferred NLT motif.^39, 40^ As both of these secondary glycosylation sites are in the lumen and ApNGT is too large to enter through the pores of spNC-4 shells, this result suggests that hexosylation occurs before cage assembly, which in turn implies that assembly must be slower than ApNGT labeling.

**Table 1.** Observed peptides with and without hexosylation and the corresponding percentage of hexosylation of spNC-4 following co-expression with ApNGT. Peak areas were extracted from ion chromatograms of the relevant m/z values. The percentage of hexosylation was calculated as the ratio of the hexosylated peptide peak area to the total peak area of the peptide species with and without hexose modification.

| Peptide sequence | Modification | m/z | Peak area | Percent hexosylation |
| --- | --- | --- | --- | --- |
| K.AANAGAGAGAMATPHFDY<sup>39</sup>NASVVSK.G | n/a | 793.38<sup>3+</sup> | 11644085325 | 52.79% |
|  | Hex | 847.40<sup>3+</sup> | 13021160482 |  |
| K.GLA<sup>49</sup>NLSLELRK.P | n/a | 607.39<sup>2+</sup> | 15526098144 | 7.01% |
|  | Hex | 688.39<sup>2+</sup> | 1170375437 |  |

**Table 2.** Observed peptides with and without hexosylation and the corresponding percentage of hexosylation of spNC-4-GI following co-expression with ApNGT. Peak areas were obtained from ion chromatograms of the respective m/z signals. The percentage of hexosylation was calculated as the ratio of the hexosylated peptide peak area to the total peak area of the peptide species with and without hexose modification. The glycosylated asparagine residue within the engineered glycotag is denoted as “g11,” reflecting its position as the eleventh residue in the tag.

| Peptide sequence | Modification | m/z | Peak area | Percent hexosylation |
| --- | --- | --- | --- | --- |
| K. AANAGAGAGAMATPHFDY<sup>39</sup>NASVVSK.G | n/a | 793.38<sup>3+</sup> | 16880619199 | 52.79% |
|  | Hex | 847.40<sup>3+</sup> | 16903654168 |  |
| K. GLA<sup>49</sup>NLSLELRK.P | n/a | 607.39<sup>2+</sup> | 23441407897 | 7.01% |
|  | Hex | 688.39<sup>2+</sup> | 1531898861 |  |
| K.SLRGTEGSGSGAHATA<sup>g11</sup>NATAHASHHHHHH | n/a | 587.47<sup>5+</sup> | 33734574 | 99.5% |
|  | Hex | 619.88<sup>5+</sup> | 7968745942 |  |

Analysis of the C subunits of spNC-4-GI and spNC-4 revealed additional chemical modifications. Some hexosylated, gluconoylated, and phosphogluconoylated species, representing approximately 10–15% of the total sample, were detected by mass spectrometry (Figure S9). Glycosylation of the C subunit most likely occurs on Asn136, which is the first residue in an NHT sequon, whereas gluconoylation and phosphogluconoylation are common nonenzymatic posttranslational modifications in *E. coli*.^41^ For applications for which these additional modifications are undesired, they can be eliminated by simple mutagenesis of the respective sites (glycosylation) or suppressed by metabolic pathway engineering (gluconoylation and phosphogluconoylation).^42^

### Targeting spNC-4 to cells using external antibody-recruiting peptides

We recently demonstrated that large, positively supercharged lumazine synthase cage variants can be targeted to specific cells with monoclonal antibodies by functionalizing their surface with antibody binding domains (ABDs).^36^ To apply this approach to spNC-4, we genetically fused a 58-amino-acid ABD derived from protein A, which binds the Fc region of antibodies with a *K*_D_ of 20 nM,^43, 44^ to the external C-terminus of the N fragment (Figure 5A). The resulting variant, spNC-4-ABD, was purified by anion-exchange chromatography (Figure S10A). TEM showed that it assembles into slightly larger cages (33.1 ± 0.9 nm) than the construct lacking the ABD (Figure 5B). Mass spectrometry and SDS-PAGE confirmed full presentation of the ABD segment on each N fragment (Figure 5C, Figure S10B-C).

**Figure 5.**
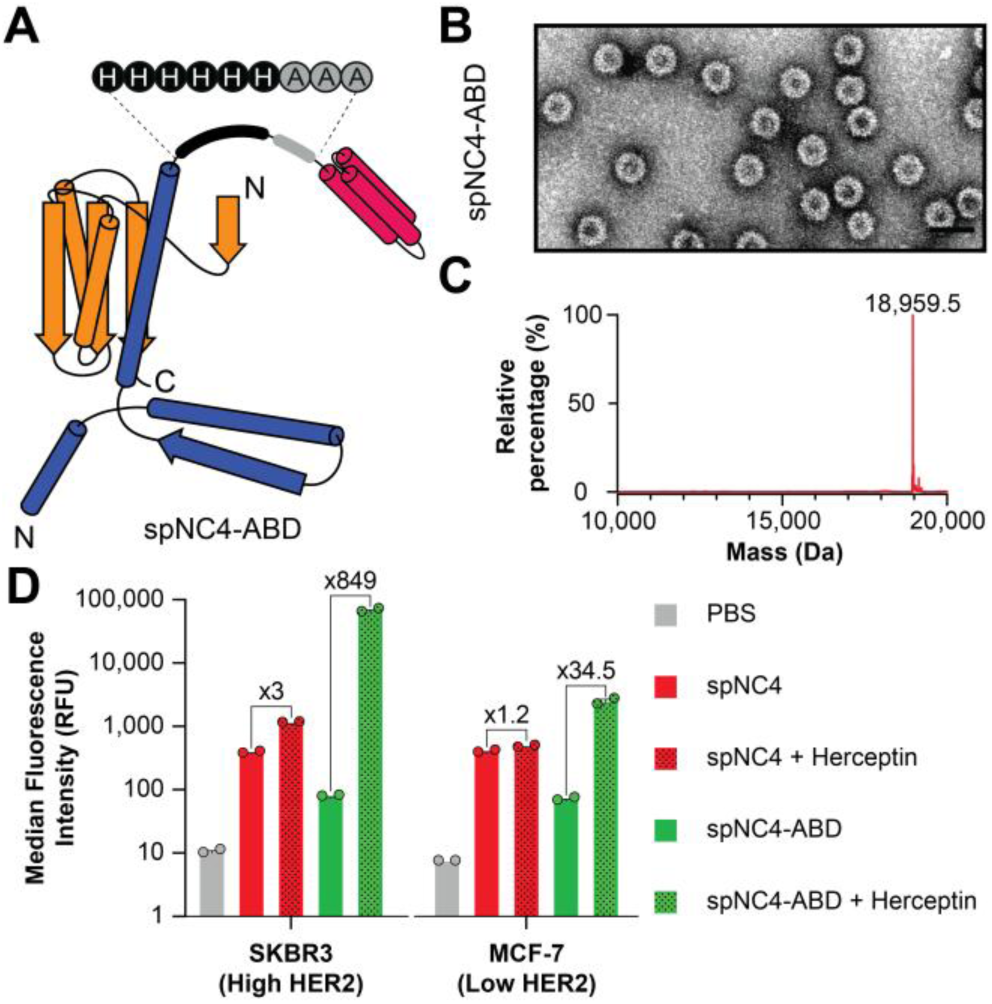
Cell targeting with antibody-binding spNC-4 cages. (**A**) Schematic representation of an spNC-4 monomer displaying an antibody-binding domain (ABD, pink). ABD peptide is connected to the capsid subunit by an eight-residue linker comprising a hexahistidine tag (black spheres) and an alanine-rich spacer (gray). (**B**) TEM and (**C**) MS analyses of purified spNC-4-ABD cages. (**D**) Cellular uptake of fluorescently labeled cages in two cell types was measured by flow cytometry. Gray bars indicate cell autofluorescence, while red and green bars show the median fluorescence intensities for labeled spNC-4 and spNC-4-ABD cages, respectively. Dotted bars denote samples incubated with the HER2-targeting antibody Herceptin. Flow cytometry data were obtained in a single experiment using duplicate wells per condition and identical protein preparations.

As cellular targets for spNC-4-ABD, we chose the human breast cancer cell lines SKBR-3 and MCF-7, which respectively have high and low expression levels of the human epidermal growth receptor 2 (HER2).^45, 46^ The monoclonal antibody trastuzumab (herceptin), which specifically recognizes HER2 and is used clinically for the treatment of metastatic breast cancer,^47^ served as the targeting antibody. The spNC-4 and spNC-4-ABD nucleocapsids were fluorescently labeled with Atto565-NHS-ester to assess cellular uptake by SKBR-3 and MCF-7, and internalization was monitored by flow cytometry in the presence and absence of herceptin. Neither spNC-4 nor spNC-4-ABD is preferentially taken up by either cell line in the absence of antibody. However, addition of herceptin increases the uptake of spNC-4-ABD by SKBR-3 cells substantially (∼850-fold) and by MCF-7 cells to a lesser extent (35-fold), reflecting the different HER2 expression levels of the two cell lines (Figure 5D). In contrast, adding the targeting antibody to the spNC-4 variant lacking the ABD tags did not improve uptake.

## CONCLUSION

*Aquifex aeolicus* lumazine synthase (AaLS) has emerged as a highly versatile scaffold for engineering protein containers. In its native form, AaLS assembles into a thermally stable, 60-subunit *T*=1 icosahedral capsid, with both N-and C-termini on its exterior surface.^24^ Through rational design and directed evolution, diverse AaLS variants have been created, including supercharged forms capable of encapsulating oppositely charged cargo, as well as novel tetrahedral and icosahedral assemblies built entirely from pentamers.^12^ Circular permutation has expanded the AaLS structural repertoire, yielding spherical and tubular architectures with inward-facing N- and C-termini.^48^ The successful splitting of the circularly permuted, 240-subunit NC-4 nucleocapsid into a two-component, 480-subunit protein nanocage with modifiable termini on both interior and exterior surfaces provides yet another demonstration of the remarkable structural plasticity and robustness of the AaLS platform.

Despite the inherent complexity of the domain-swapped NC-4 scaffold, the two spNC-4 fragments efficiently co-assemble in the cytosol of *E. coli*, likely aided by co-translation from a bicistronic mRNA, similar to what has been observed for Splitcore, an engineered nanoparticle obtained by splitting the capsid protein of the hepatitis B virus.^20^ Structural analyses by cryo-EM and solid-state NMR confirm that the assembled structure closely mirrors the parental icosahedral *T*=4 capsid, with only localized perturbations near the newly engineered termini. The C subunit forms the structural core of the shell wall, while the N subunit comprises a single shell-spanning helix, the helix-strand element responsible for the domain swap in NC-4, as well as the interior-facing RNA-binding λN+ peptide and an exterior-facing hexahistidine purification tag.

Functionally, spNC-4 retains the ability of NC-4 to package and protect its encoding mRNA, though with somewhat lower efficiency and specificity, encapsulating one rather than two or three full-length mRNAs per particle. This decrease may reflect the greater complexity of two-component assembly or altered RNA folding caused by the 76-nt insert separating the open reading frames, which could disrupt the packaging cassette that we evolved for optimal capsid assembly and RNA encapsidation. Further rounds of evolution or synonymous recoding of the bicistronic transcript could conceivably restore full packaging efficiency.

The new N- and C-termini in spNC-4 provide convenient handles for functionalizing the capsid exterior. For example, displaying a 17-amino acid peptide containing glycosylation motifs enabled site-specific glycosylation when the split nucleocapsid was co-produced with a glucosyltransferase. In principle, these starter monosaccharides could be elaborated in *E. coli* using established glyco-engineering pathways to produce capsids with more complex glycan structures.^37, 38^ Such constructs would be useful for investigating lectin-sugar interactions, enhancing cellular uptake, triggering glycan-dependent immune responses, and modulating immunogenicity.

Similarly, fusing a 58-amino-acid antibody-binding domain to the nanocage surface facilitated recruitment of targeting immunoglobulins and cell-specific uptake of the encapsulated cargo. Genetically appending nanobodies or computationally designed minibinders that recognize desired cell-surface receptors should also be feasible,^49–51^ provided the fused domains can be produced in soluble form and are compatible in size with the capsid architecture. Such constructs could be used as general and evolvable display vehicles,^52^ as long as the bicistronic mRNA encoding the fusion protein does not exceed the ∼2500 nt packaging capacity of the capsid. Encapsulation intrinsically couples genotype and phenotype, thereby enabling genetic selection of displayed protein variants for tasks such as targeted delivery or receptor-specific binding.

The ability to rationally design and precisely assemble large protein nanostructures capable of encapsulating a diverse range of biologically relevant cargo while displaying multiple functional elements on both interior and exterior surfaces makes the spNC-4 system attractive for applications in transfection, drug delivery, and vaccine development. Creating split versions of other lumazine synthase-derived molecular containers, which have been shown to accommodate proteins,^53^ nucleic acids,^54^ and imaging agents,^55^ would further expand their utility across biotechnology and synthetic biology.

## ASSOCIATED CONTENT

*Supporting Information Available*: This material is available free of charge via the Internet. Experimental procedures and additional figures and data (PDF).

## AUTHOR INFORMATION

### Notes

The authors declare no competing financial interests.

## ACKNOWLEDGMENTS

We thank the Scientific Center for Optical and Electron Microscopy (ScopeM) of ETH Zurich for help with cryo-EM Data collection and elaboration, and Thomas Wiegand and Alexander A. Malär for help with solid-state NMR experiments. This work was supported by the ETH Zurich. N.T. is grateful for Human Frontier Science Program Long-term Fellowship (LT000426/2015-L). A.S. is grateful for a Marie Skłodowska-Curie Individual Fellowship (LEVERAGE mRNA).

## Supplementary information

### SUPPLEMENTARY TABLES

**Table S1.**
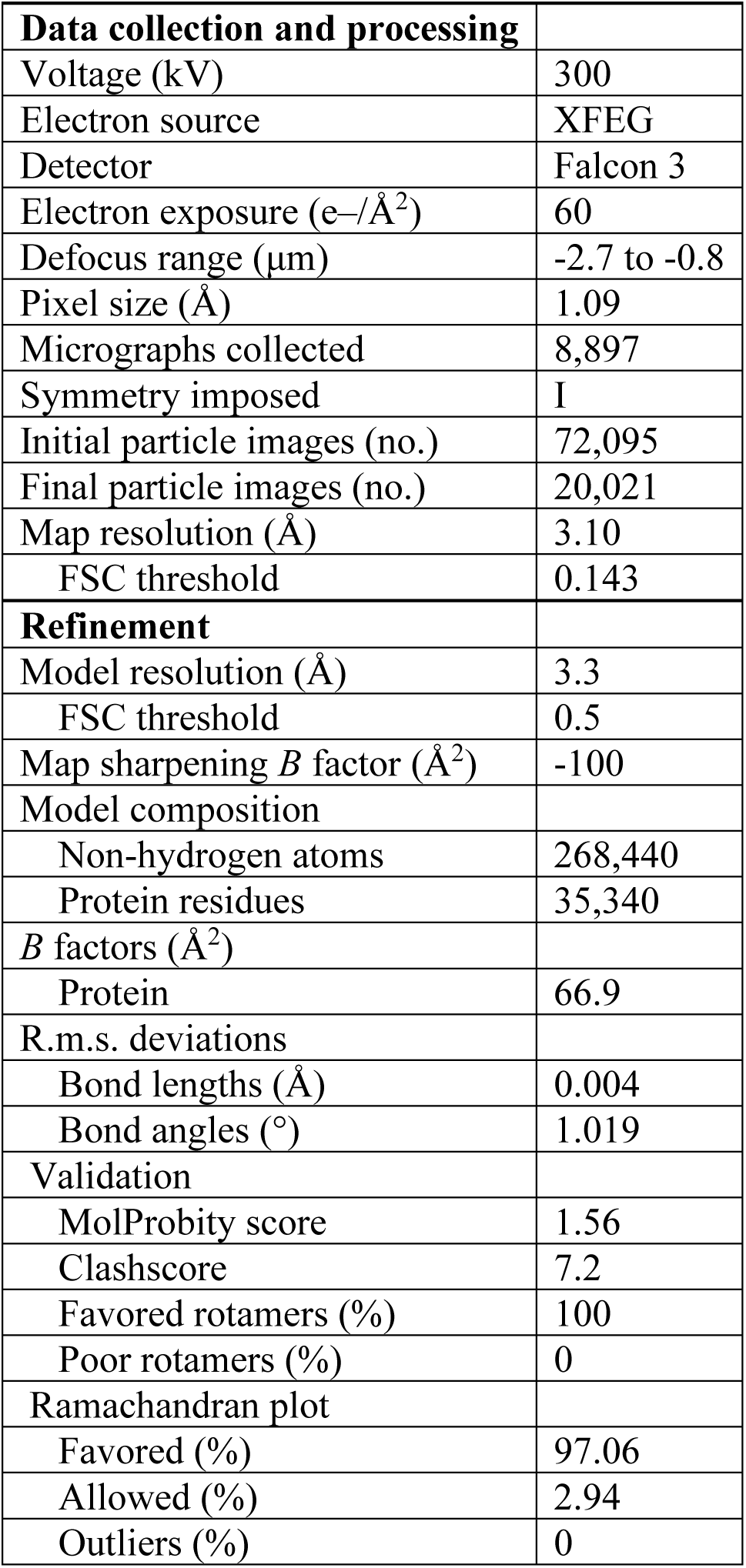
Cryo-EM data.

|  |  |
| --- | --- |
| **Data collection and processing** |  |
| Voltage (kV) | 300 |
| Electron source | XFEG |
| Detector | Falcon 3 |
| Electron exposure (e-/Å<sup>2</sup>) | 60 |
| Defocus range (μm) | -2.7 to -0.8 |
| Pixel size (Å) | 1.09 |
| Micrographs collected | 8,897 |
| Symmetry imposed | I |
| Initial particle images (no.) | 72,095 |
| Final particle images (no.) | 20,021 |
| Map resolution (Å) | 3.10 |
| FSC threshold | 0.143 |
| **Refinement** |  |
| Model resolution (Å) | 3.3 |
| FSC threshold | 0.5 |
| Map sharpening *B* factor (Å<sup>2</sup>) | -100 |
| Model composition |  |
| Non-hydrogen atoms | 268,440 |
| Protein residues | 35,340 |
| *B* factors (Å<sup>2</sup>) |  |
| Protein | 66.9 |
| R.m.s. deviations |  |
| Bond lengths (Å) | 0.004 |
| Bond angles (°) | 1.019 |
| Validation |  |
| MolProbity score | 1.56 |
| Clashscore | 7.2 |
| Favored rotamers (%) | 100 |
| Poor rotamers (%) | 0 |
| Ramachandran plot |  |
| Favored (%) | 97.06 |
| Allowed (%) | 2.94 |
| Outliers (%) | 0 |

**Table S2.** 2D and 3D NMR spectra for resonance assignment. Lowercase letters in the experiment names indicate magnetization transfers where no corresponding dimension was recorded.

| spNC-4 | NC-4 |
| --- | --- |
| **carbon-detected** |  |
| $^{13}\text{C}$-$^{13}\text{C}$ DARR | $^{13}\text{C}$-$^{13}\text{C}$ DARR |
| NCA, NCO | NCA, NCO |
| NCACX | NCACX |
| NCACB, CANCO, NCOCX | - |
| CANcoCA, NcoCACB | - |
| **proton-detected** |  |
| hNH, hCH | hNH, hCH |
| hCANH | hCANH |
| hNCAH, hNcoCAH, hCAcoNH | - |

**Table S3.** Experimental parameters for solid-state NMR experiments with spNC-4.

|  | **DARR** | **NCA** | **NCO** |
| --- | --- | --- | --- |
| MAS frequency/ kHz | 17 | 17 | 17 |
| Field/ T | 20 | 20 | 20 |
| Number of scans | 8 | 8 | 8 |
| t1 increment | 3072 | 1600 | 1600 |
| SW (t1)/ ppm | 467.7 | 773.7 | 773.7 |
| Acqu. time (t1)/ ms | 15.4 | 12 | 12 |
| t2 increments | 3072 | 3072 | 3072 |
| SW (t2)/ ppm | 467.7 | 467.7 | 467.7 |
| Acqu. time (t2)/ ms | 15.4 | 15.4 | 15.4 |
| <sup>1</sup>H decoupling/ kHz | SPINAL64 / 90 | SPINAL64 / 90 | SPINAL64 / 90 |
| Inter-scan delay/ s | 2.7 | 3 | 3 |
| Experiment time | 19 h | 11 h | 11 h |
| **Transfer 1** | **HC-CP** | **HN-CP** | **HN-CP** |
| Field/ kHz | 60 (1H) / 41 (13C) | 60.4 (1H) / 40 (15N) | 60.4 (1H) / 40 (15N) |
| Shape | Tangent 1H | Tangent 1H | Tangent 1H |
| Carrier/ ppm | 100 (13C) | 120 (15N) | 120 (15N) |
| Time/ ms | 0.5 | 0.7 | 0.7 |
| **Transfer 2** | **DARR** | **NCA-CP** | **NCO-CP** |
| Field/ kHz | 17 (1H) | 9.0 (15N) / 6.0 (13C) | 9.9 (15N) / 6.0 (13C) |
| Shape | - | Tangent 13C | Tangent 13C |
| Carrier/ ppm | 100 (13C) | 54.8 (13C) | 174.1 (13C) |
| Time/ ms | 20 | 6 | 3.5 |
| Window function | QSINE SSB 2.5 | QSINE SSB 3 | QSINE SSB 3 |

|  | NCACX | NCOCX | CANCO |
| --- | --- | --- | --- |
| MAS frequency/ kHz | 17 | 17 | 17 |
| Field/ T | 20 | 20 | 20 |
| Number of scans | 16 | 32 | 16 |
| t1 increment | 68 | 60 | 120 |
| SW (t1)/ ppm | 55 | 55 | 37 |
| Acqu. time (t1)/ ms | 7.2 | 6.3 | 7.6 |
| t2 increments | 128 | 112 | 60 |
| SW (t2)/ ppm | 37 | 32 | 55 |
| Acqu. time (t2)/ ms | 8.1 | 8.2 | 6.3 |
| t3 increments | 2304 | 2304 | 2304 |
| SW (t3)/ ppm | 467.7 | 467.7 | 467.7 |
| Acqu. time (t3)/ ms | 11.5 | 11.5 | 11.5 |
| 1H decoupling/ kHz | SPINAL64 / 90 | SPINAL64 / 90 | SPINAL64 / 90 |
| Interscan delay / s | 3 | 3 | 3 |
| Experiment time | 5 d | 6 d 17 h | 4 d |
| **Transfer 1** | **HN-CP** | **HN-CP** | **HCA-CP** |
| Field/ kHz | 60.4 (1H) / 40 (15N) | 60.4 (1H) / 40 (15N) | 60.4 (1H) / 42.4 (13C) |
| Shape | Tangent 1H | Tangent 1H | Tangent 1H |
| Carrier/ ppm | 120 (15N) | 120 (15N) | 54.8 (13C) |
| Time/ ms | 0.7 | 0.7 | 0.45 |
| **Transfer 2** | **NCA-CP** | **NCO-CP** | **CAN-CP** |
| Field/ kHz | 9.0 (15N) / 6.0 (13C) | 9.9 (15N) / 6.0 (13C) | 6.0 (13C) / 9.5 (15N) |
| Shape | Tangent 13C | Tangent 13C | Tangent 13C |
| Carrier/ ppm | 54.8 (13C) | 174.1 (13C) | 54.8 (13C) |
| Time/ ms | 6 | 3.5 | 6 |
| **Transfer 3** | **DARR** | **DARR** | **NCO-CP** |
| Field/ kHz | 17 (1H) | 17 (1H) | 9.7 (15N) / 6.0 (13C) |
| Shape | - | - | Tangent 13C |
| Carrier/ ppm | 174.1 (13C) | 174.1 (13C) | 174.1 (13C) |
| Time/ ms | 70 | 70 | 3.5 |
| Window function | QSINE SSB 2.5 | QSINE SSB 2.5 | QSINE SSB 2.5 |

|  | NCACB | CANcoCA | NcoCACB |
| --- | --- | --- | --- |
| MAS frequency/ kHz | 17 | 17 | 17 |
| Field/ T | 20 | 20 | 20 |
| Number of scans | 8 | 24 | 16 |
| t1 increment | 90 | 122 | 68 |
| SW (t1)/ ppm | 60 | 35 | 54 |
| Acqu. time (t1)/ ms | 8.7 | 8.2 | 7.3 |
| t2 increments | 144 | 60 | 114 |
| SW (t2)/ ppm | 42 | 55 | 34 |
| Acqu. time (t2)/ ms | 8 | 6.3 | 7.8 |
| t3 increments | 2304 | 2304 | 3072 |
| SW (t3)/ ppm | 467.7 | 467 | 467.7 |
| Acqu. time (t3)/ ms | 11.5 | 11.5 | 15.4 |
| 1H decoupling/ kHz | SPINAL64 / 90 | SPINAL64 / 90 | SPINAL64 / 90 |
| Interscan delay/ s | 3 | 3.2 | 3.2 |
| Experiment time | 3 d 15 h | 6 d 14 h | 4 d 16 h |
| **Transfer 1** | **HN-CP** | **HCA-CP** | **HN-CP** |
| Field/ kHz | 60 (1H) / 46 (15N) | 60 (1H) / 40.4 (13C) | 60 (1H) / 46 (15N) |
| Shape | Tangent 1H | Tangent 1H | Tangent 1H |
| Carrier/ ppm | 120 (15N) | 54.8 (13C) | 120 (15N) |
| Time/ ms | 0.85 | 0.4 | 0.8 |
| **Transfer 2** | **NCA-CP** | **CAN-CP** | **NCO-CP** |
| Field/ kHz | 10.9 (15N) / 6.0 (13C) | 6 (13C) / 10.1 (15N) | 20.9 (15N) / 6.0 (13C) |
| Shape | Tangent 13C | Tangent 13C | Tangent 13C |
| Carrier/ ppm | 59.3 (13C) | 54.8 (13C) | 177.2 (13C) |
| Time/ ms | 8.5 | 6 | 6 |
| **Transfer 3** | **DREAM** | **NCO-CP** | **Mod. band-selective CP** |
| Field/ kHz | 7.0 (13C) | 10.6 (15N) / 6.0 (13C) | 7.2 (13C) |
| Shape | Tangent 13C | Tangent 13C | Tangent 13C |
| Carrier/ ppm | 56.7 (13C) | 174.1 (13C) | 177.2 (13C) |
| Time/ ms | 1.1 | 3.5 | 7.5 |
| **Transfer 4** | - | **Mod. band-selective CP** | **DREAM** |
| Field/ kHz | - | 7.3 (13C) | 7.3 (13C) |
| Shape | - | Tangent 13C | Tangent 13C |
| Carrier/ ppm | - | 174.1 (13C) | 55.1 (13C) |
| Time/ ms | - | 5.6 | 1.2 |
| Window function | QSINE SSB 3.0 | QSINE SSB 2.5 | QSINE SSB 2.5 |

|  | hNH | hCH | hCANH |
| --- | --- | --- | --- |
| MAS frequency/ kHz | 100 | 100 | 100 |
| Field/ T | 20 | 20 | 20 |
| Number of scans | 64 | 80 | 24 |
| t1 increment | 800 | 800 | 104 |
| SW (t1)/ ppm | 180 | 200 | 32 |
| Acqu. time (t1)/ ms | 25.8 | 9.4 | 7.6 |
| t2 increments | 2048 | 2048 | 48 |
| SW (t2)/ ppm | 46.7 | 46.7 | 38 |
| Acqu. time (t2)/ ms | 25.8 | 25.8 | 7.3 |
| t3 increments | - | - | 2048 |
| SW (t3)/ ppm | - | - | 46.7 |
| Acqu. time (t3)/ ms | - | - | 25.8 |
| 1H decoupling/ kHz | swfTPPM / 10.2 | swfTPPM / 9.8 | swfTPPM / 9.6 |
| 15N decoupling/ kHz | WALTZ64 / 5.4 | - | WALTZ64 / 5.3 |
| 13C decoupling/ kHz | - | WALTZ64 / 5.4 | WALTZ64 / 5.0 |
| Water suppression/ kHz | MISS. 120 ms / 20.3 | MISS. 120 ms / 19.7 | MISS. 120 ms / 19.3 |
| Interscan delay/ s | 1.2 | 1 | 1 |
| Experiment time | 19 h | 20 h | 1 d 15 h |
| **Transfer 1** | **HN-CP** | **HC-CP** | **HC-CP** |
| Field/ kHz | 76.3 (1H) / 16.1 (15N) | 74.6 (1H) / 16.7 (13C) | 76.9 (1H) / 16.5 (13C) |
| Shape | Tangent 1H | Tangent 1H | Tangent 1H |
| Carrier/ ppm | 117.5 (15N) | 56.0 (13C) | 56.0 (13C) |
| Time/ ms | 1.1 | 0.5 | 0.5 |
| **Transfer 2** | **NH-CP** | **CH-CP** | **CN-CP** |
| Field/ kHz | 16.1 (15N) / 76.3 (1H) | 16.7 (13C) / 74.6 (1H) | 65.1 (13C) / 32.3 (15N) |
| Shape | Tangent 1H | Tangent 1H | Tangent 13C |
| Carrier/ ppm | 4.8 (1H) | 4.8 (1H) | 117.5 |
| Time/ ms | 0.8 | 0.6 | 17 |
| **Transfer 3** | - | - | **NH-CP** |
| Field/ kHz | - | - | 15.9 (15N) / 77.3 (1H) |
| Shape | - | - | Tangent 1H |
| Carrier/ ppm | - | - | 4.8 (1H) |
| Time/ ms | - | - | 1.2 |
| Window function | QSINE SSB 3.0 | QSINE SSB 3.0 | QSINE SSB 2.5 |
| TDeff | TD2eff 1024 | TD2eff 1024 | TD3eff 1024 |

|  | hNCAH | hNcoCAH | hCAcoNH |
| --- | --- | --- | --- |
| MAS frequency/ kHz | 100 | 100 | 100 |
| Field/ T | 20 | 20 | 20 |
| Number of scans | 28 | 72 | 52 |
| t1 increment | 44 | 46 | 108 |
| SW (t1)/ ppm | 38 | 38 | 32 |
| Acqu. time (t1)/ ms | 6.7 | 7 | 7.9 |
| t2 increments | 96 | 98 | 48 |
| SW (t2)/ ppm | 32 | 32 | 38 |
| Acqu. time (t2)/ ms | 7 | 7.2 | 7.3 |
| t3 increments | 2048 | 2048 | 2048 |
| SW (t3)/ ppm | 46.7 | 46.7 | 46.7 |
| Acqu. time (t3)/ ms | 25.8 | 25.8 | 25.8 |
| 1H decoupling/ kHz | swfTPPM / 9.8 | swfTPPM / 9.4 | swfTPPM / 9.4 |
| 15N decoupling/ kHz | WALTZ64 / 5.4 | WALTZ64 / 4.8 | WALTZ64 / 5.5 |
| 13C decoupling/ kHz | WALTZ64 / 5.5 | WALTZ64 / 4.69 | WALTZ64 / 5.5 |
| Water suppression/ kHz | MISS. 120 ms / 19.6 | MISS. 120 ms / 18.7 | MISS. 120 ms / 18.7 |
| Inter-scan delay/ s | 1 | 1 | 1 |
| Experiment time | 1 d 15 h | 4 d 11 h | 3 d 16 h |
| **Transfer 1** | **HN-CP** | **HN-CP** | **HC-CP** |
| Field/ kHz | 78.2 (1H) / 15.7 (15N) | 78.3 (1H) / 15.6 (15N) | 76.9 (1H) / 16.7 (13C) |
| Shape | Tangent 1H | Tangent 1H | Tangent 1H |
| Carrier/ ppm | 117.5 (15N) | 117.5 (15N) | 51.8 (13C) |
| Time/ ms | 0.9 | 1 | 0.5 |
| **Transfer 2** | **NC-CP** | **NCO-CP** | **DREAM** |
| Field/ kHz | 31.5 (15N) / 66.9 (13C) | 31.7 (15N) / 63.2 (13C) | 45.1 (13C) |
| Shape | Tangent 13C | Tangent 13C | Tangent 13C |
| Carrier/ ppm | 56.0 (13C) | 222.1 (13C) | 184.2 (13C) |
| Time/ ms | 19 | 14 | 7 |
| **Transfer 3** | **CH-CP** | **DREAM** | **CN-CP** |
| Field/ kHz | 16.7 (13C) / 75.0 (1H) | 46.1 (13C) | 63.8 (13C) / 32.3 (15N) |
| Shape | Tangent 1H | Tangent 13C | Tangent 13C |
| Carrier/ ppm | 4.8 (1H) | 156.1 (13C) | 117.5 |
| Time/ ms | 0.65 | 9 | 14.5 |
| **Transfer 4** | - | **CAH-CP** | **NH-CP** |
| Field/ kHz | - | 15.6 (13C) / 75.0 (1H) | 15.9 (15N) / 74.2 (1H) |
| Shape | - | Tangent 1H | Tangent 1H |
| Carrier/ ppm | - | 4.8 (1H) | 4.8 (1H) |
| Time/ ms | - | 0.4 | 1.2 |
| Window function | QSINE SSB 2.5 | QSINE SSB 2.5 | QSINE SSB 2.7 |
| TDeff | TD3eff 1024 | TD3eff 1024 | TD3eff 1024 |

**Table S4.**
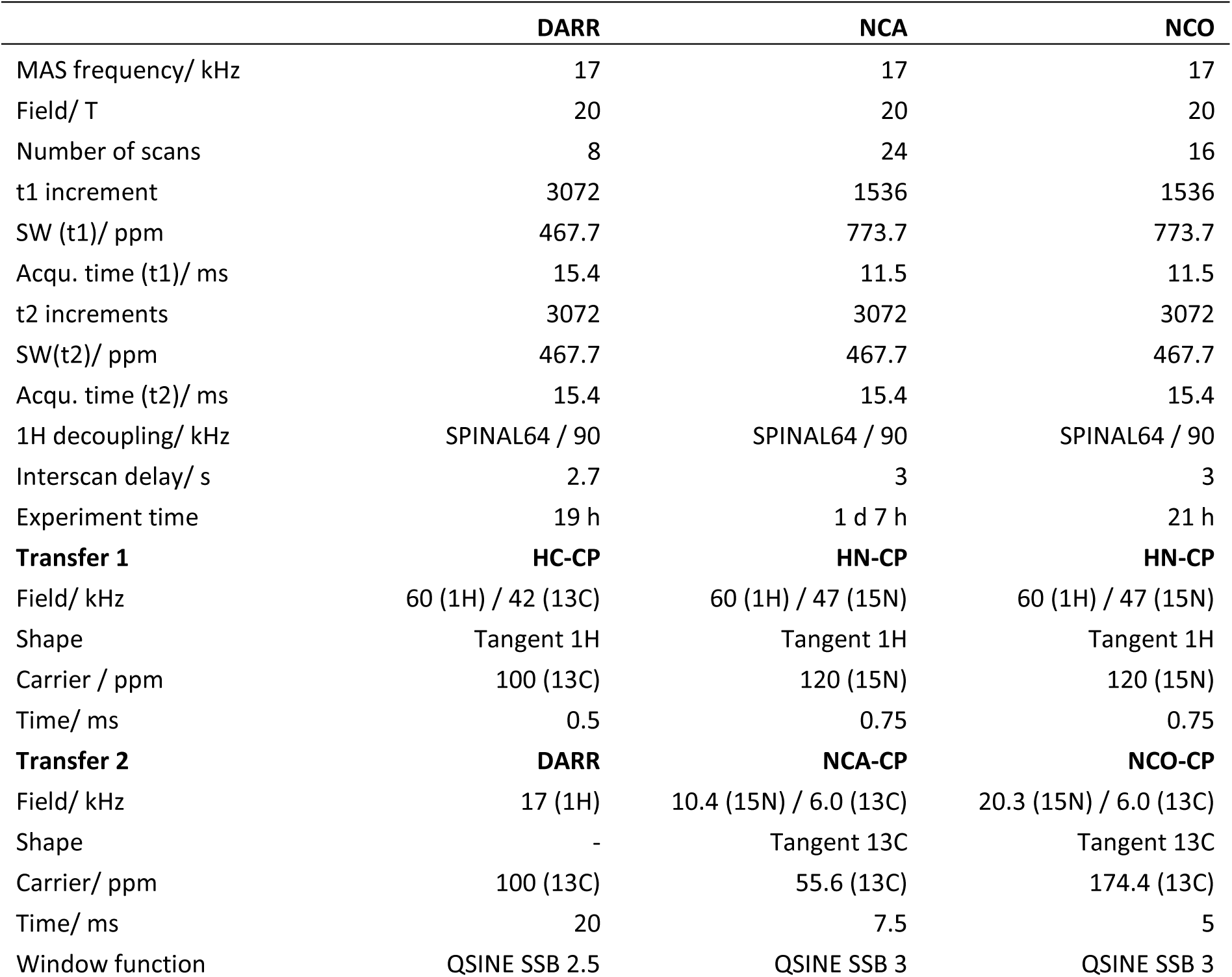
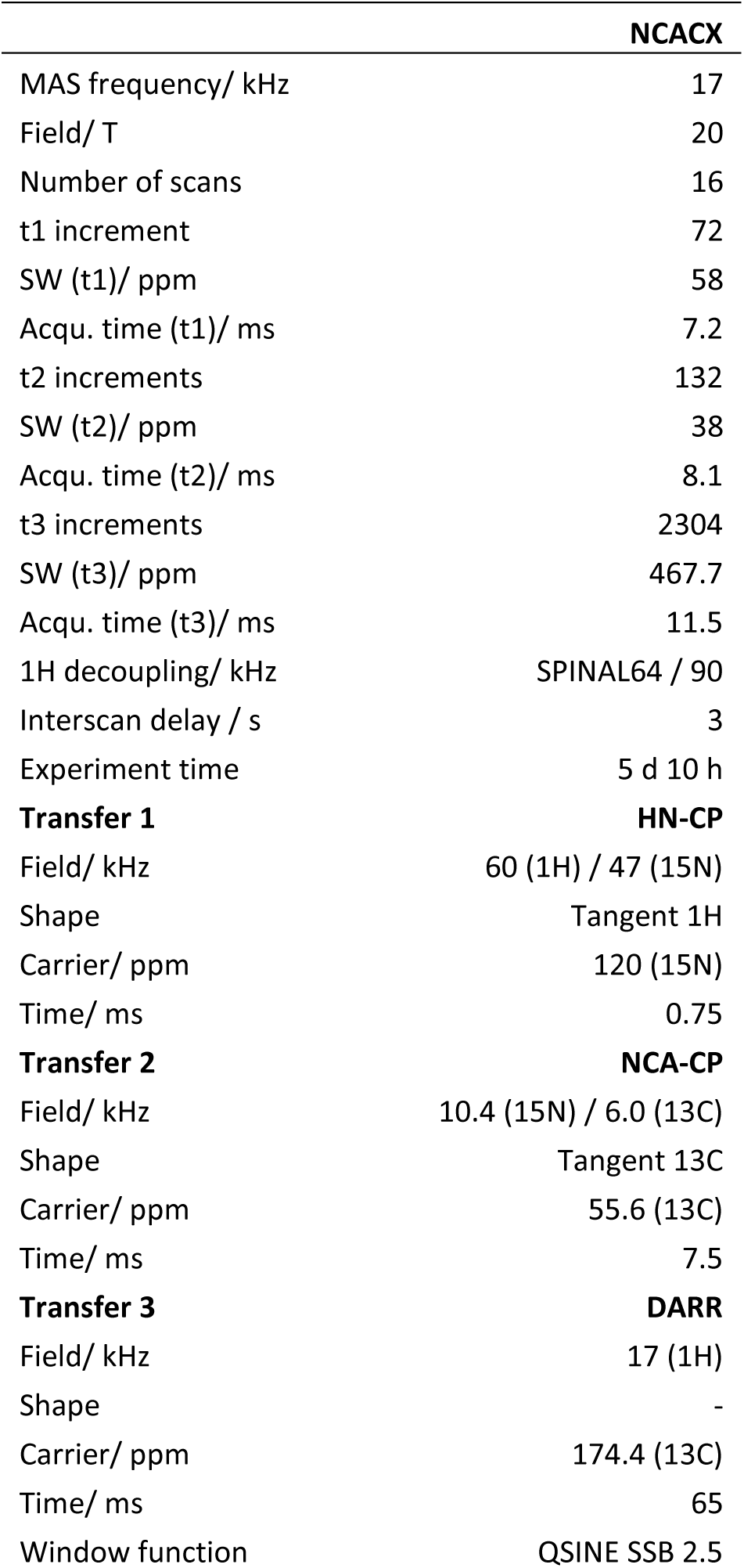
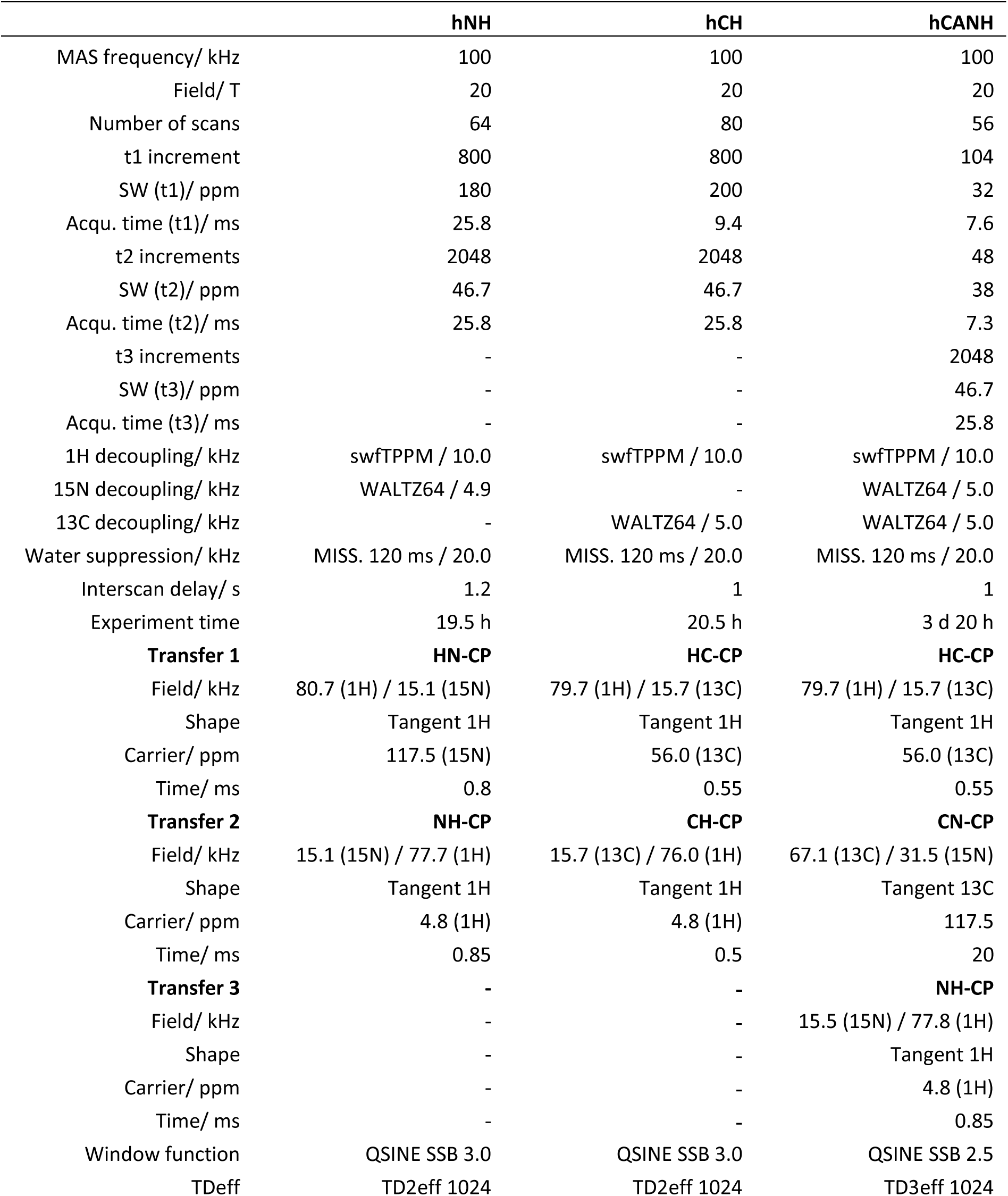
Experimental parameters for solid-state NMR experiments with NC-4.

|  | **DARR** | **NCA** | **NCO** |
| --- | --- | --- | --- |
| MAS frequency/ kHz | 17 | 17 | 17 |
| Field/ T | 20 | 20 | 20 |
| Number of scans | 8 | 24 | 16 |
| t1 increment | 3072 | 1536 | 1536 |
| SW (t1)/ ppm | 467.7 | 773.7 | 773.7 |
| Acqu. time (t1)/ ms | 15.4 | 11.5 | 11.5 |
| t2 increments | 3072 | 3072 | 3072 |
| SW(t2)/ ppm | 467.7 | 467.7 | 467.7 |
| Acqu. time (t2)/ ms | 15.4 | 15.4 | 15.4 |
| <sup>1</sup>H decoupling/ kHz | SPINAL64 / 90 | SPINAL64 / 90 | SPINAL64 / 90 |
| Inter-scan delay/ s | 2.7 | 3 | 3 |
| Experiment time | 19 h | 1 d 7 h | 21 h |
| **Transfer 1** | **HC-CP** | **HN-CP** | **HN-CP** |
| Field/ kHz | 60 (1H) / 42 (13C) | 60 (1H) / 47 (15N) | 60 (1H) / 47 (15N) |
| Shape | Tangent 1H | Tangent 1H | Tangent 1H |
| Carrier / ppm | 100 (13C) | 120 (15N) | 120 (15N) |
| Time/ ms | 0.5 | 0.75 | 0.75 |
| **Transfer 2** | **DARR** | **NCA-CP** | **NCO-CP** |
| Field/ kHz | 17 (1H) | 10.4 (15N) / 6.0 (13C) | 20.3 (15N) / 6.0 (13C) |
| Shape | - | Tangent 13C | Tangent 13C |
| Carrier/ ppm | 100 (13C) | 55.6 (13C) | 174.4 (13C) |
| Time/ ms | 20 | 7.5 | 5 |
| Window function | QSINE SSB 2.5 | QSINE SSB 3 | QSINE SSB 3 |

|  | NCACX |
| --- | --- |
| MAS frequency/ kHz | 17 |
| Field/ T | 20 |
| Number of scans | 16 |
| t1 increment | 72 |
| SW (t1)/ ppm | 58 |
| Acqu. time (t1)/ ms | 7.2 |
| t2 increments | 132 |
| SW (t2)/ ppm | 38 |
| Acqu. time (t2)/ ms | 8.1 |
| t3 increments | 2304 |
| SW (t3)/ ppm | 467.7 |
| Acqu. time (t3)/ ms | 11.5 |
| 1H decoupling/ kHz | SPINAL64 / 90 |
| Interscan delay / s | 3 |
| Experiment time | 5 d 10 h |
| **Transfer 1** | **HN-CP** |
| Field/ kHz | 60 (1H) / 47 (15N) |
| Shape | Tangent 1H |
| Carrier/ ppm | 120 (15N) |
| Time/ ms | 0.75 |
| **Transfer 2** | **NCA-CP** |
| Field/ kHz | 10.4 (15N) / 6.0 (13C) |
| Shape | Tangent 13C |
| Carrier/ ppm | 55.6 (13C) |
| Time/ ms | 7.5 |
| **Transfer 3** | **DARR** |
| Field/ kHz | 17 (1H) |
| Shape | - |
| Carrier/ ppm | 174.4 (13C) |
| Time/ ms | 65 |
| Window function | QSINE SSB 2.5 |

|  | hNH | hCH | hCANH |
| --- | --- | --- | --- |
| MAS frequency/ kHz | 100 | 100 | 100 |
| Field/ T | 20 | 20 | 20 |
| Number of scans | 64 | 80 | 56 |
| t1 increment | 800 | 800 | 104 |
| SW (t1)/ ppm | 180 | 200 | 32 |
| Acqu. time (t1)/ ms | 25.8 | 9.4 | 7.6 |
| t2 increments | 2048 | 2048 | 48 |
| SW (t2)/ ppm | 46.7 | 46.7 | 38 |
| Acqu. time (t2)/ ms | 25.8 | 25.8 | 7.3 |
| t3 increments | - | - | 2048 |
| SW (t3)/ ppm | - | - | 46.7 |
| Acqu. time (t3)/ ms | - | - | 25.8 |
| 1H decoupling/ kHz | swfTPPM / 10.0 | swfTPPM / 10.0 | swfTPPM / 10.0 |
| 15N decoupling/ kHz | WALTZ64 / 4.9 | - | WALTZ64 / 5.0 |
| 13C decoupling/ kHz | - | WALTZ64 / 5.0 | WALTZ64 / 5.0 |
| Water suppression/ kHz | MISS. 120 ms / 20.0 | MISS. 120 ms / 20.0 | MISS. 120 ms / 20.0 |
| Interscan delay/ s | 1.2 | 1 | 1 |
| Experiment time | 19.5 h | 20.5 h | 3 d 20 h |
| **Transfer 1** | **HN-CP** | **HC-CP** | **HC-CP** |
| Field/ kHz | 80.7 (1H) / 15.1 (15N) | 79.7 (1H) / 15.7 (13C) | 79.7 (1H) / 15.7 (13C) |
| Shape | Tangent 1H | Tangent 1H | Tangent 1H |
| Carrier/ ppm | 117.5 (15N) | 56.0 (13C) | 56.0 (13C) |
| Time/ ms | 0.8 | 0.55 | 0.55 |
| **Transfer 2** | **NH-CP** | **CH-CP** | **CN-CP** |
| Field/ kHz | 15.1 (15N) / 77.7 (1H) | 15.7 (13C) / 76.0 (1H) | 67.1 (13C) / 31.5 (15N) |
| Shape | Tangent 1H | Tangent 1H | Tangent 13C |
| Carrier/ ppm | 4.8 (1H) | 4.8 (1H) | 117.5 |
| Time/ ms | 0.85 | 0.5 | 20 |
| **Transfer 3** | - | - | **NH-CP** |
| Field/ kHz | - | - | 15.5 (15N) / 77.8 (1H) |
| Shape | - | - | Tangent 1H |
| Carrier/ ppm | - | - | 4.8 (1H) |
| Time/ ms | - | - | 0.85 |
| Window function | QSINE SSB 3.0 | QSINE SSB 3.0 | QSINE SSB 2.5 |
| TDeff | TD2eff 1024 | TD2eff 1024 | TD3eff 1024 |

**Table S5.** NMR assignment completeness for spNC-4 and NC-4. (**A**) The percentage of assigned atoms considering all residues in the full-length proteins, and (**B**) the percentage calculated excluding the first 38 amino acids of the flexible N-terminus.

| atom type | **(A)** full-length |  | **(B)** without N-terminus |  |
| --- | --- | --- | --- | --- |
|  | spNC-4 (%) | NC-4 (%) | spNC-4 (%) | NC-4 (%) |
| backbone | 53 | 50 | 66 | 61 |
| sidechain non-H | 35 | 32 | 43 | 40 |
| H | 43 | 37 | 53 | 45 |
| HA | 44 | - | 54 | - |

### SUPPLEMENTARY FIGURES

**Figure S1.**
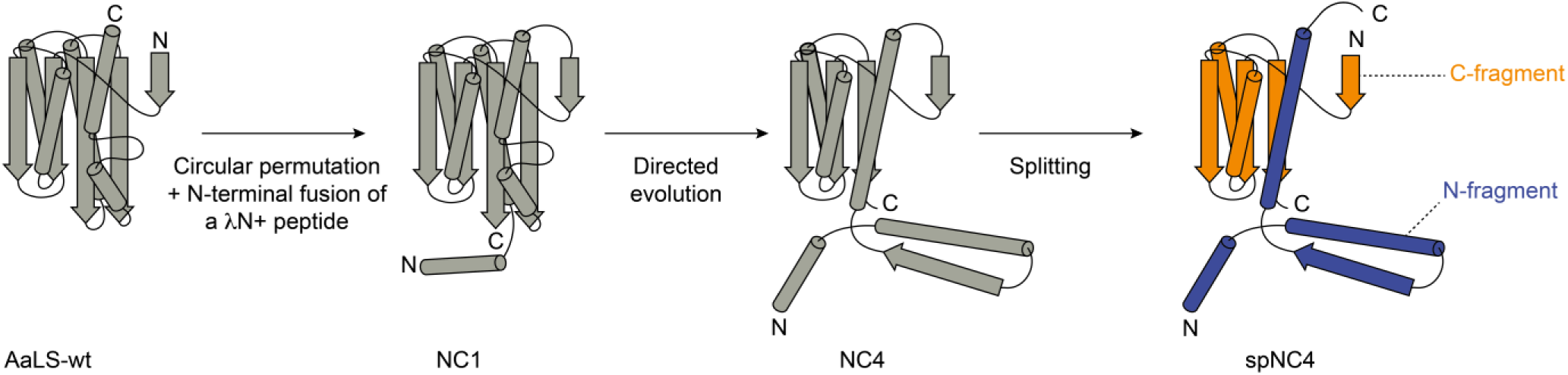
Simplified models illustrating changes in protein topology. The N and C fragments of spNC-4 are shown in blue and orange, respectively, for clear differentiation.

**Figure S2.**
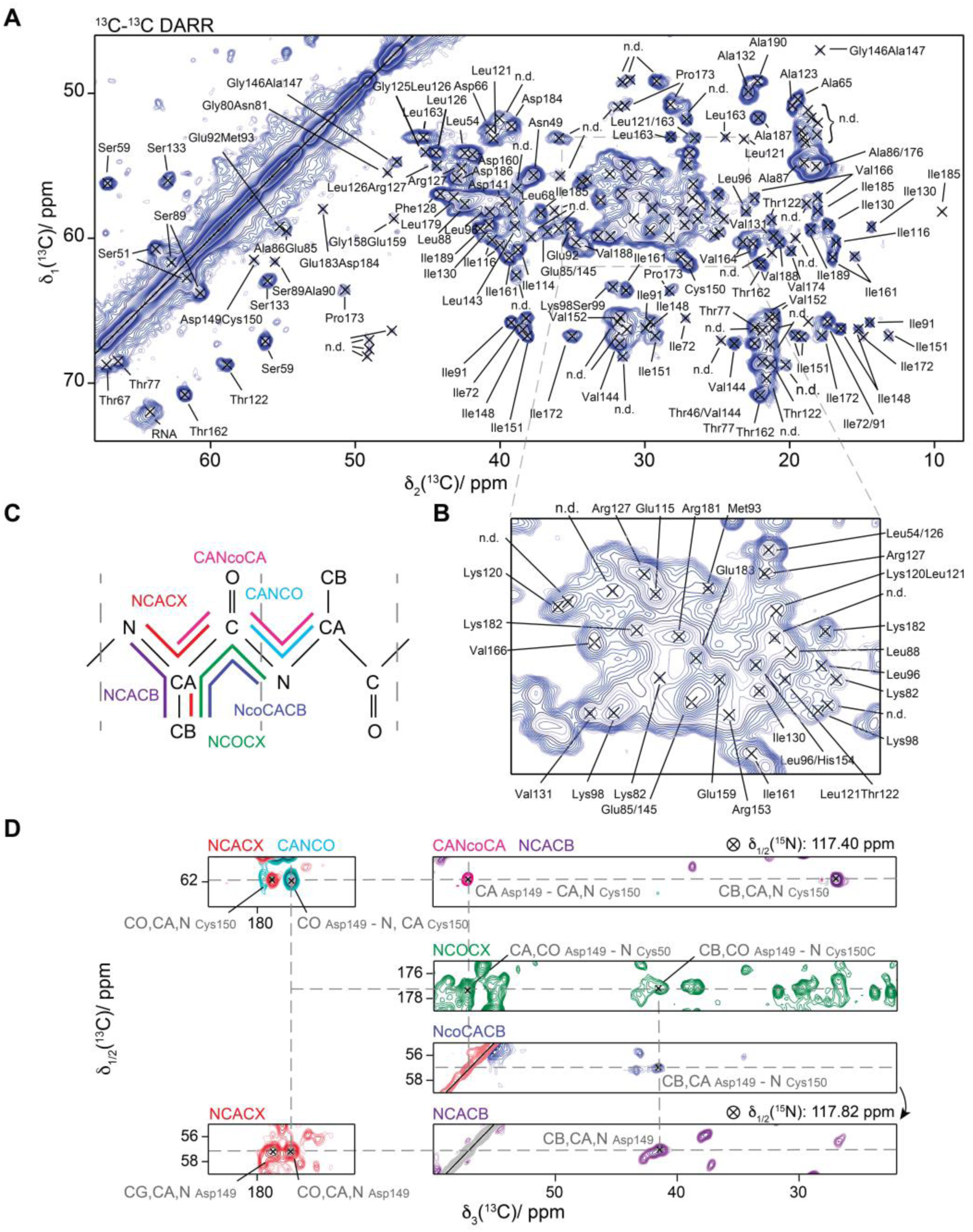
Assignment of spNC-4 with carbon-detected solid-state NMR experiments. (**A**) ^13^C-^13^C DARR spectrum recorded at 850 MHz with 17 kHz MAS in a 3.2 mm rotor; ‘n.d.’ indicates ambiguous assignments. (A) Expanded view of the crowded spectral region. (**C**) Assignments were performed by backbone walk using NCACB, NCACX, NCOCX, CANCO, CANcoCA, and NcoCACB 3D experiments. (**D**) Example of the assignment procedure, highlighting three-dimensional spectra showing N, CO, CA, and CB resonances for Cys150, and N, CO, CA, CB, and CG resonances for Asp149.

**Figure S3.**
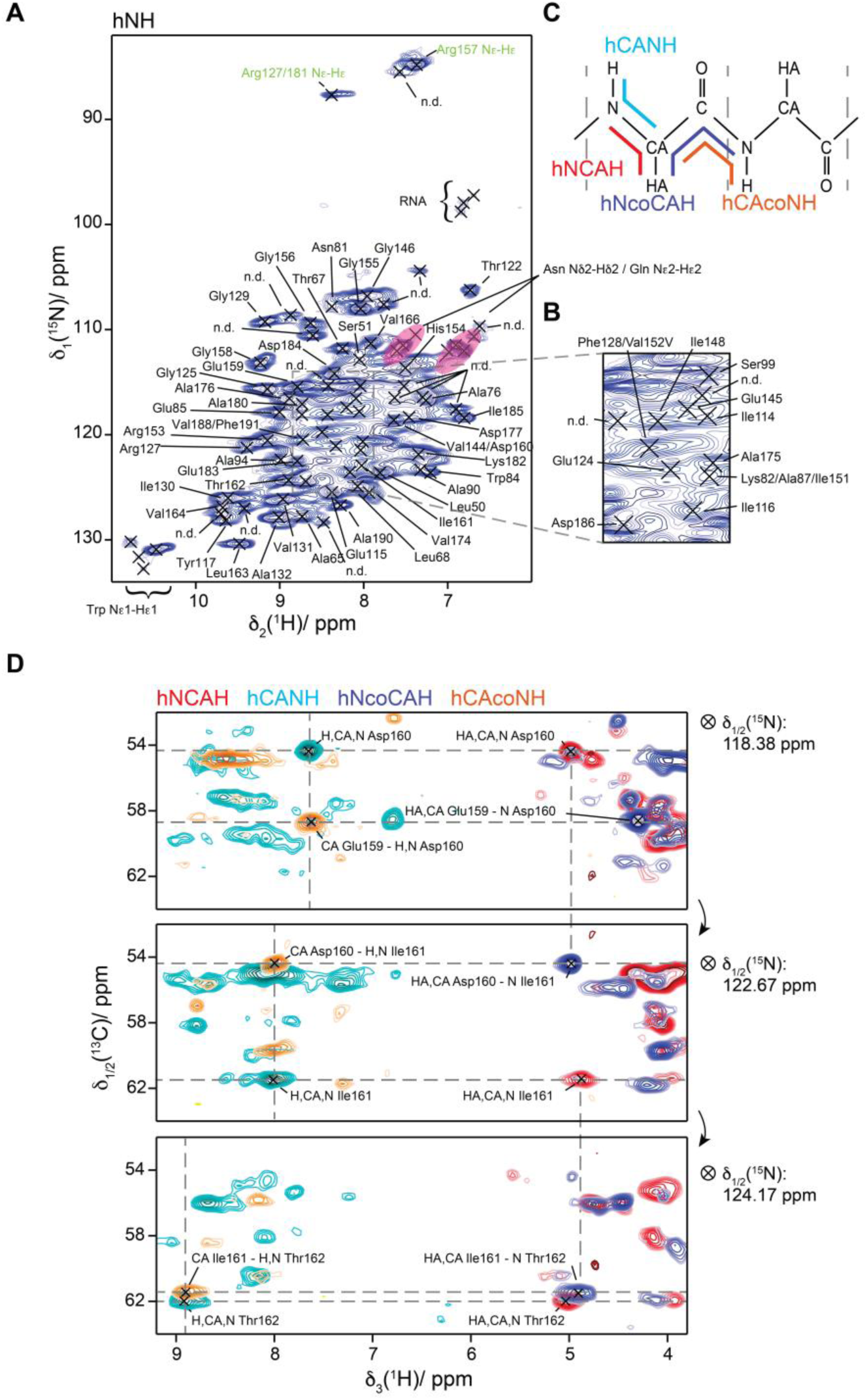
Assignment of amide and HA protons in spNC-4 by solid-state NMR. (**A**) hNH two-dimensional spectrum with assignments, acquired at 850 MHz using a 0.7 mm rotor and 100 kHz MAS. Green labels indicate tentative assignments where sequential connectivity could not be established. Asparagine (Asn) and glutamine (Gln) sidechains could not be unambiguously assigned; corresponding peaks are highlighted in pink. ‘n.d.’ denotes additional ambiguous peaks. (**B**) Expanded view of the hNH region, illustrating peak overlap and annotations. (**C**) Magnetization pathways for the hNCAH, hCANH, hNcoCAH, and hCAcoNH experiments. The sequential assignment strategy with these spectra is shown in panel (**D**), exemplified by Asp160, Ile161, and Thr162. Nitrogen and carbon resonance assignments from carbon-detected spectra (Figure S2) facilitated the assignment of amide and HA protons.

**Figure S4.**
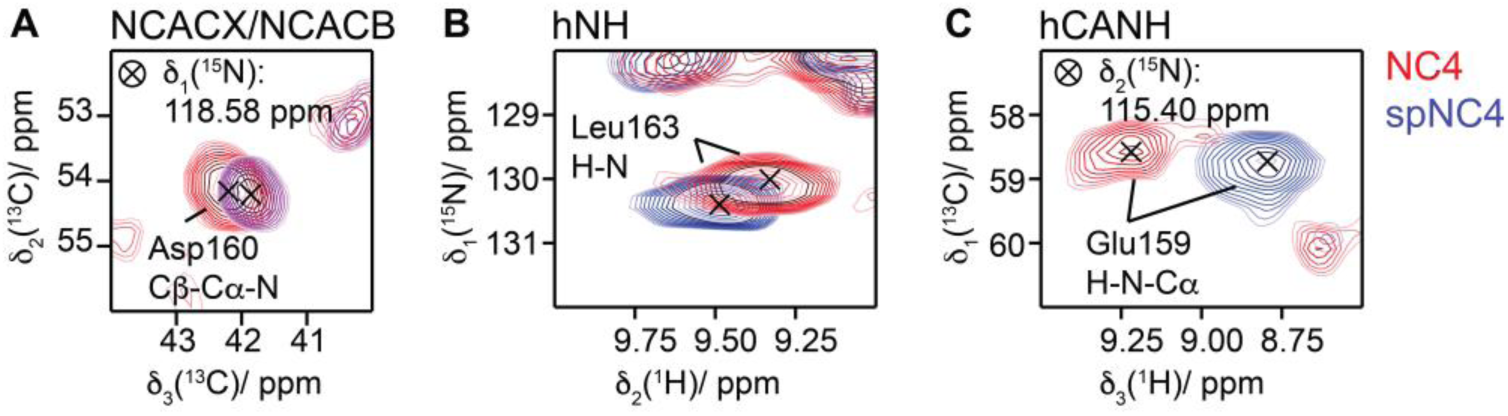
Chemical shift differences between NC-4 and spNC-4: residues 159-163. (**A**) Chemical shift of the Asp160 peak in the NCACX spectrum of NC-4 (red) compared with the NCACB spectrum of spNC-4 (purple). (**B**) hNH spectra of spNC-4 (blue) and NC-4 (red) showing the chemical shift difference in the Leu163 amide proton resonance. (**C**) hCANH spectra of spNC-4 (blue) and NC-4 (red) illustrating chemical shift changes in the Glu159 amide proton resonance.

**Figure S5.**
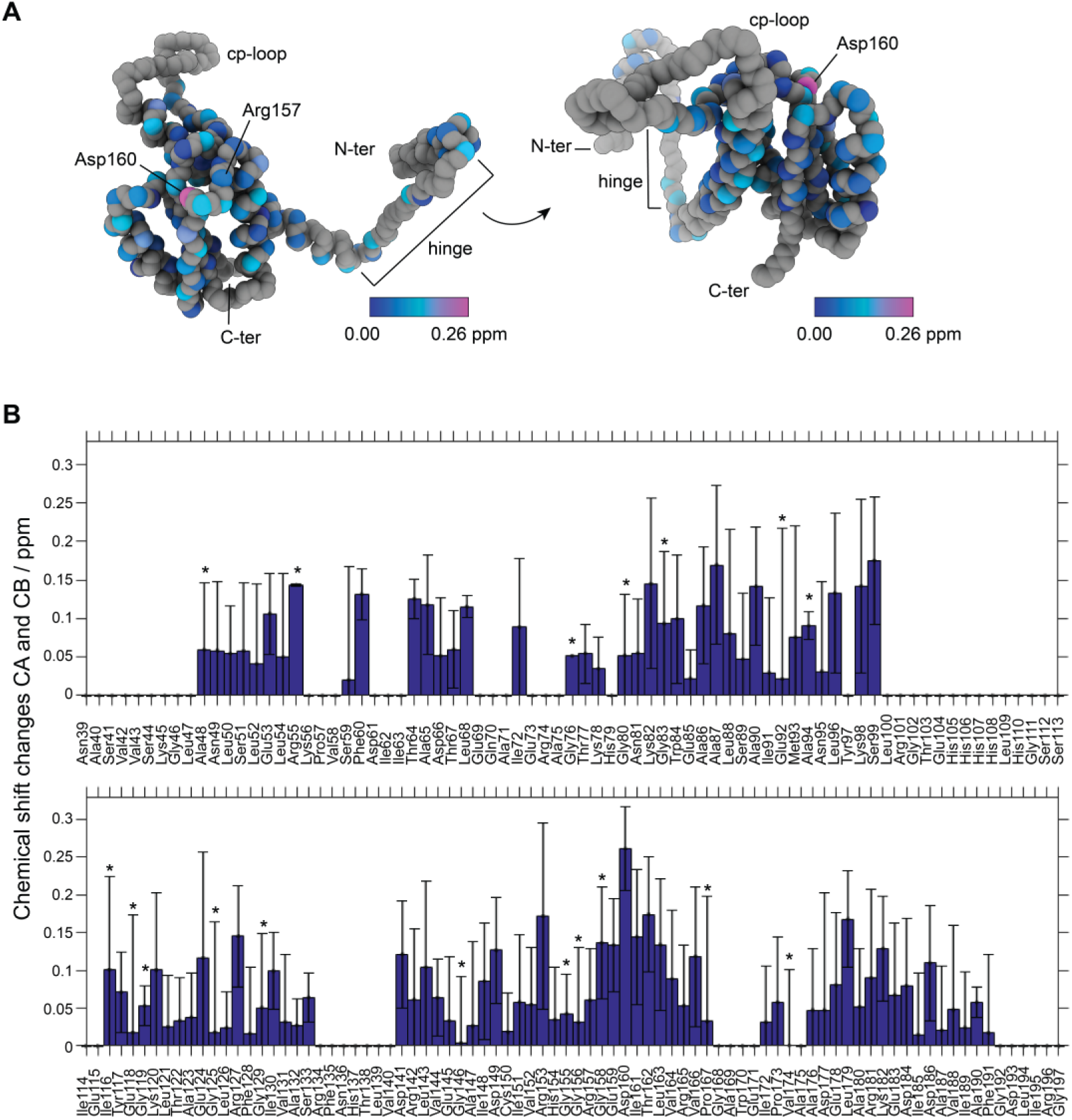
CA and CB chemical shift differences between NC-4 and spNC-4. (**A**) NC-4 structure (pdb code: 7A4J^1^) showing CA, C’, and N atoms as gray spheres. For assigned CA and CB atoms, the chemical shift differences between NC-4 and spNC-4, calculated with the formula 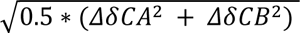, are mapped onto the NC-4 structure at the CA positions using a color scale (blue=0., pink=0.26 ppm); Asp160 shows the largest shift (0.26 ppm). (**B**) Residue-specific chemical shift differences for CA and CB atoms. Error bars indicate ± σ, calculated by error propagation of the standard deviations from peak assignments in multiple spectra using CcpNmr version 2.^2^ Stars denote residues for which only the CA resonance was assigned; for these, only CA chemical shifts are shown.

**Figure S6.**
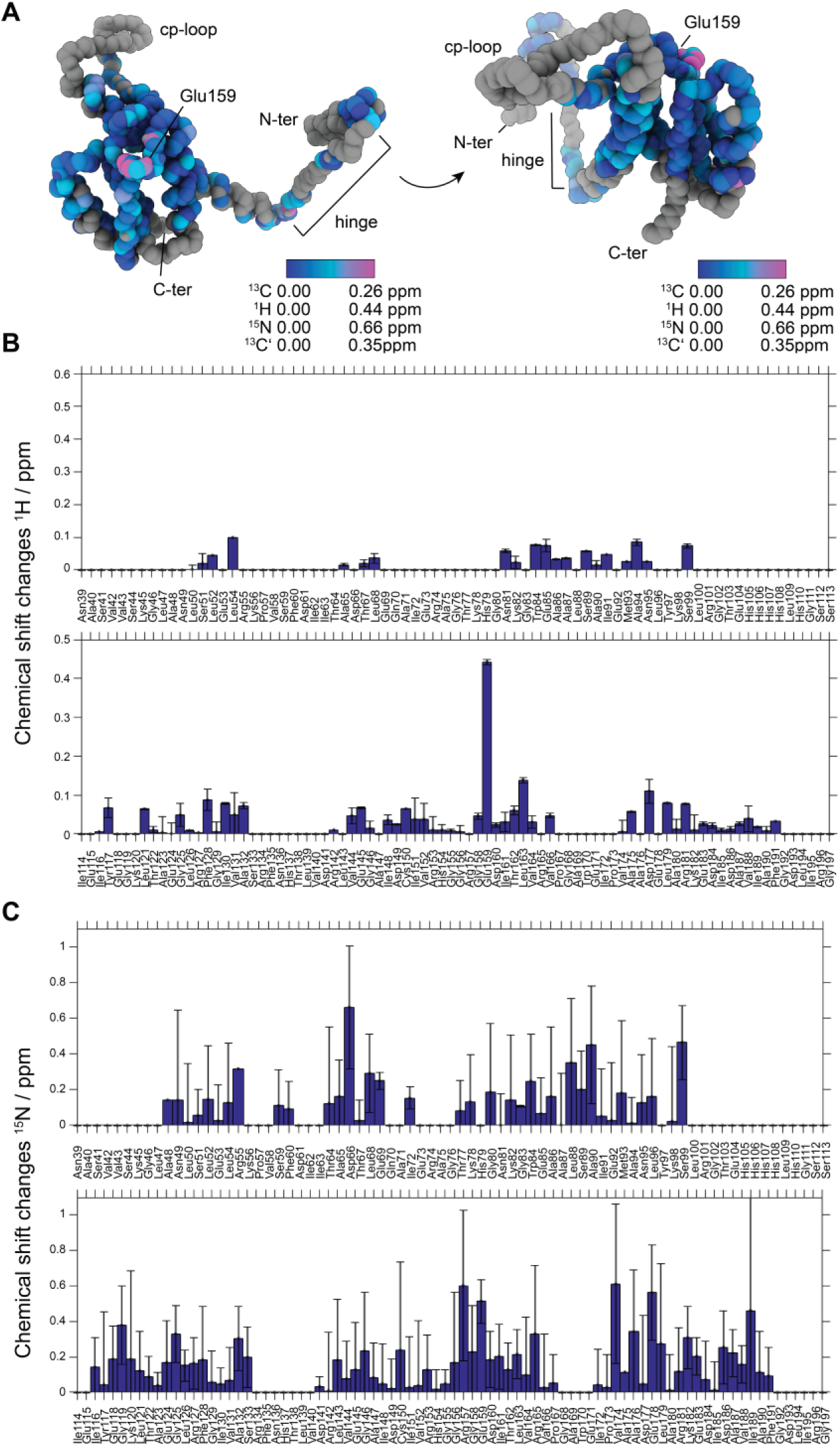
^1^H and ^15^N chemical shift changes between NC-4 and spNC-4. (**A**) NC-4 structure (pdb code: 7A4J^1^) showing CA, C’, H, and N atoms as gray spheres. Chemical shift differences for assigned CA and CB carbons, as well as H, N and C’ atoms, are mapped using color scales: blue = 0.0 and pink = 0.26 ppm for carbon atoms; 0.44 ppm for hydrogens; and 0.66 ppm for nitrogens. Glu159 exhibits the largest amide proton shift (0.44 ppm). Chemical shift changes of C’ atoms are presented separately as a bar plot in Figure S7. (**B**) Residue-specific chemical shift differences for amide protons, with error bars representing ± σ, calculated by error propagation from standard deviations of peak assignments across multiple spectra analyzed in CcpNmr version 2.^2^ (**C**) Residue-specific chemical shift differences for nitrogen atoms, with error bars similarly determined.

**Figure S7.**
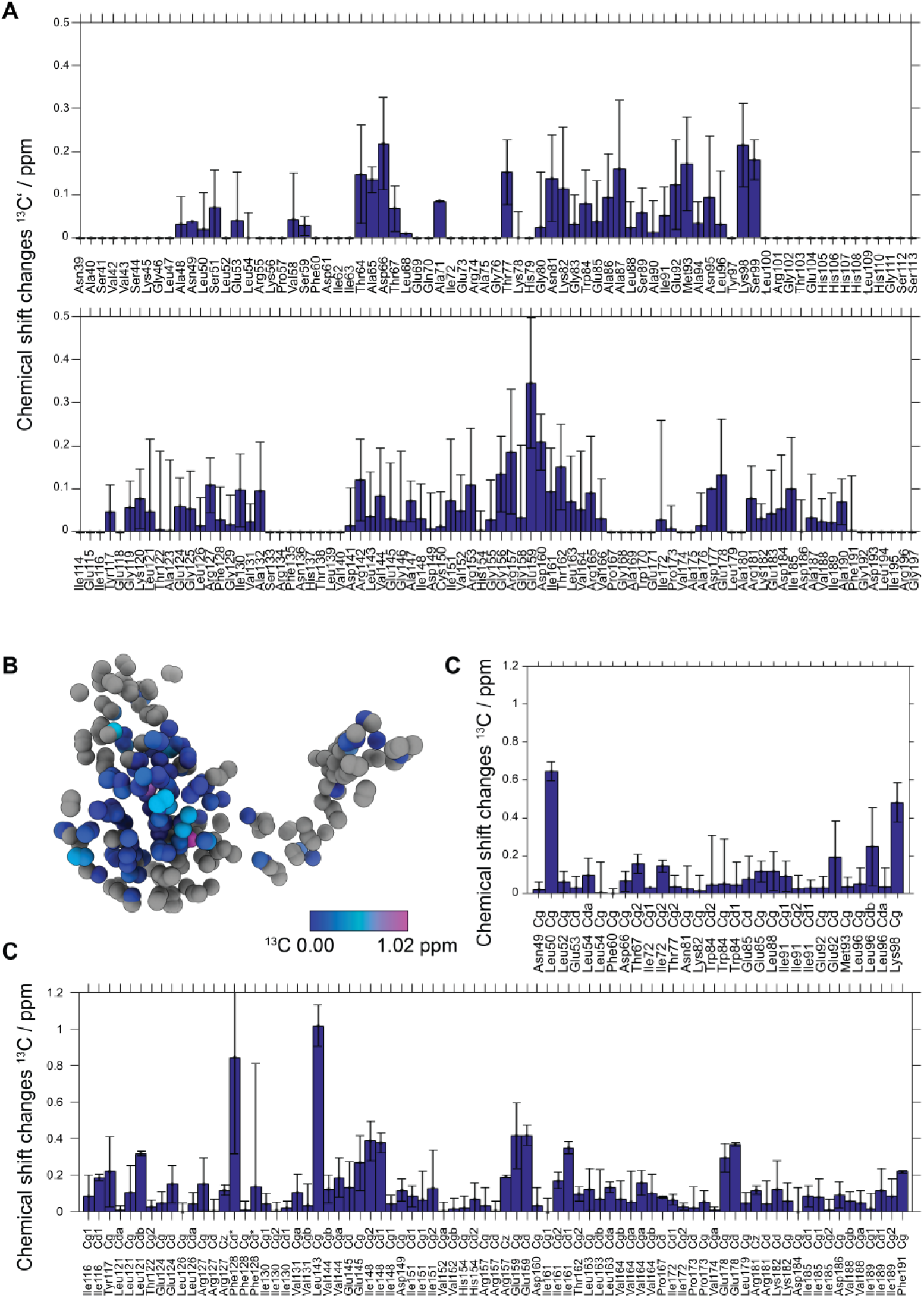
Chemical shift differences between NC-4 and spNC-4. (**A**) Chemical shift differences of C’ atoms between NC-4 and spNC-4. (**B**) NC-4 structure (pdb code: 7A4J^1^) showing Cd, Cg, and Cz sidechain atoms colored according to chemical shift differences. Phe128 Cd and Leu143 Cg exhibit the largest shifts. (B) Residue-specific chemical shift differences of sidechain carbon atoms are presented as a bar plot. Error bars denote ± σ, calculated by error propagation of the standard deviations from multiple peak assignments.

**Figure S8.**
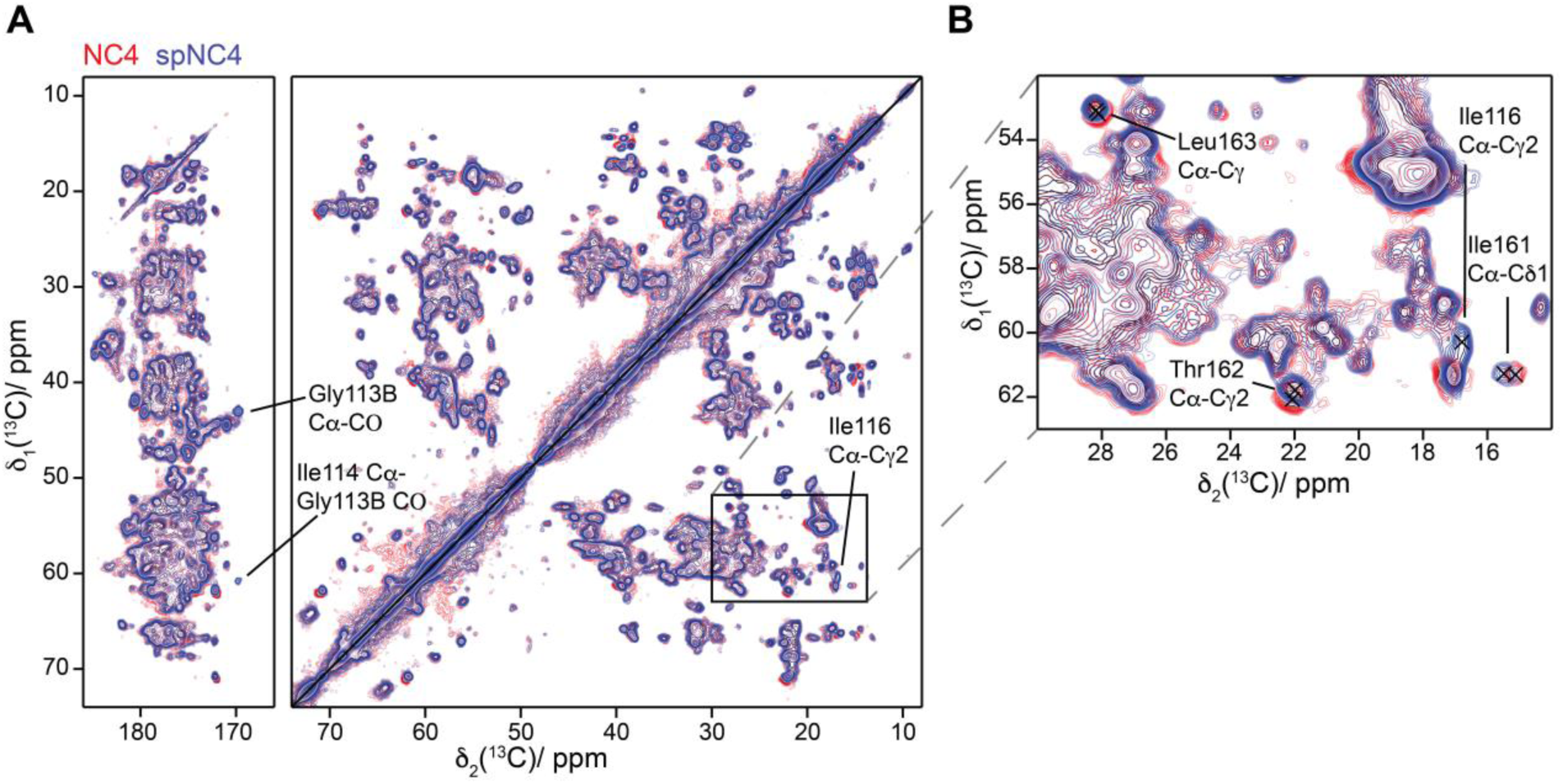
Superposition of the ^13^C-^13^C DARR spectra for spNC-4 (blue) and NC-4 (red). Spectra were acquired at 850 MHz, 17 kHz MAS, using a 3.2 mm rotor for 20 ms DARR mixing time. Residue-specific signals are colored according to their parent structure: NC-4 residues are in red, spNC-4 in blue.

**Figure S8a.**
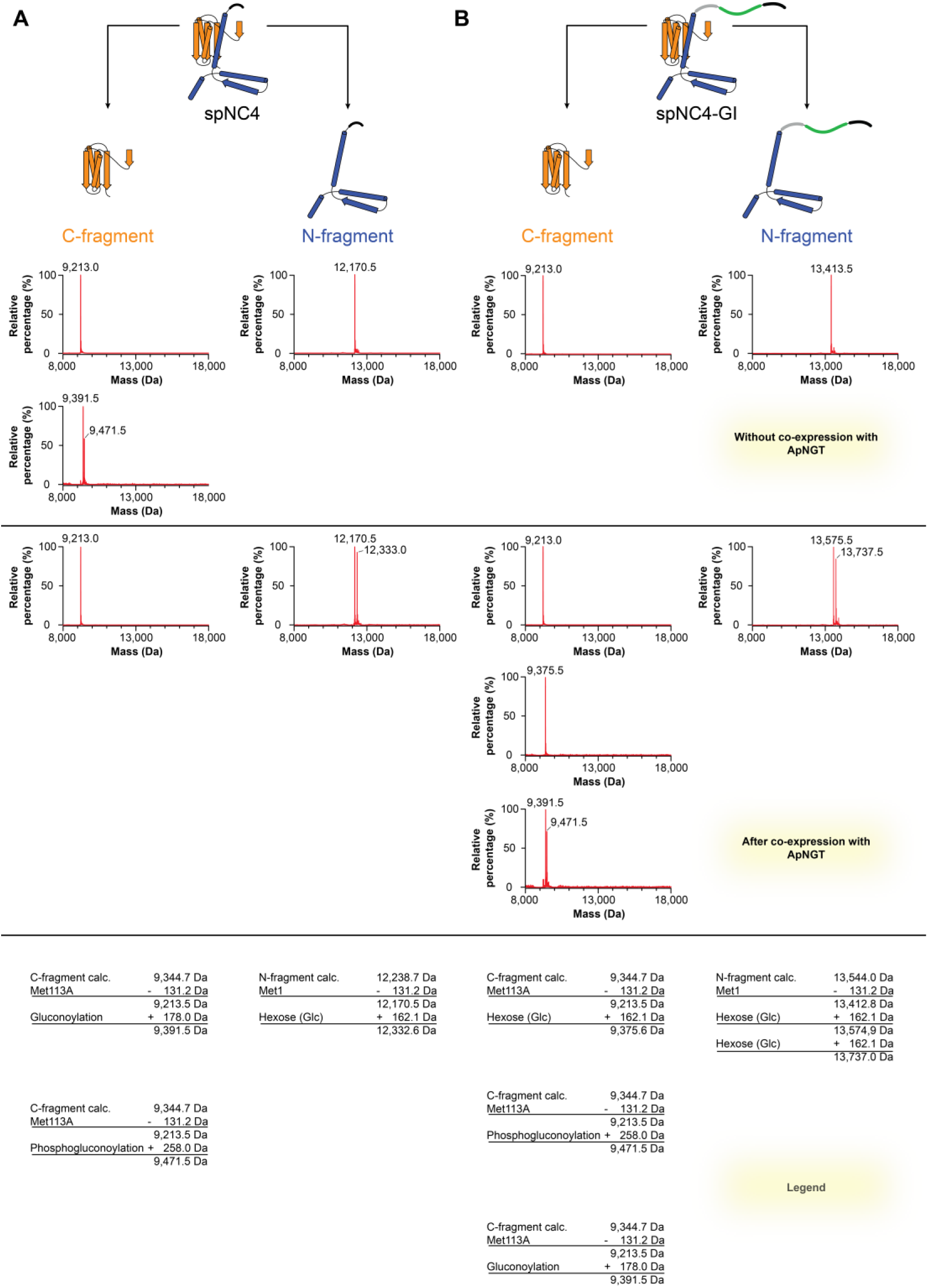
Glycosylation of spNC-4 and spNC-4-GI. MS analysis of the N and C fragments of spNC-4 (left) and spNC-4-GI (right) in the absence (top) and presence (middle) of co-expressed ApNGT. Spectra of all detected species are shown.

**Figure S9.**
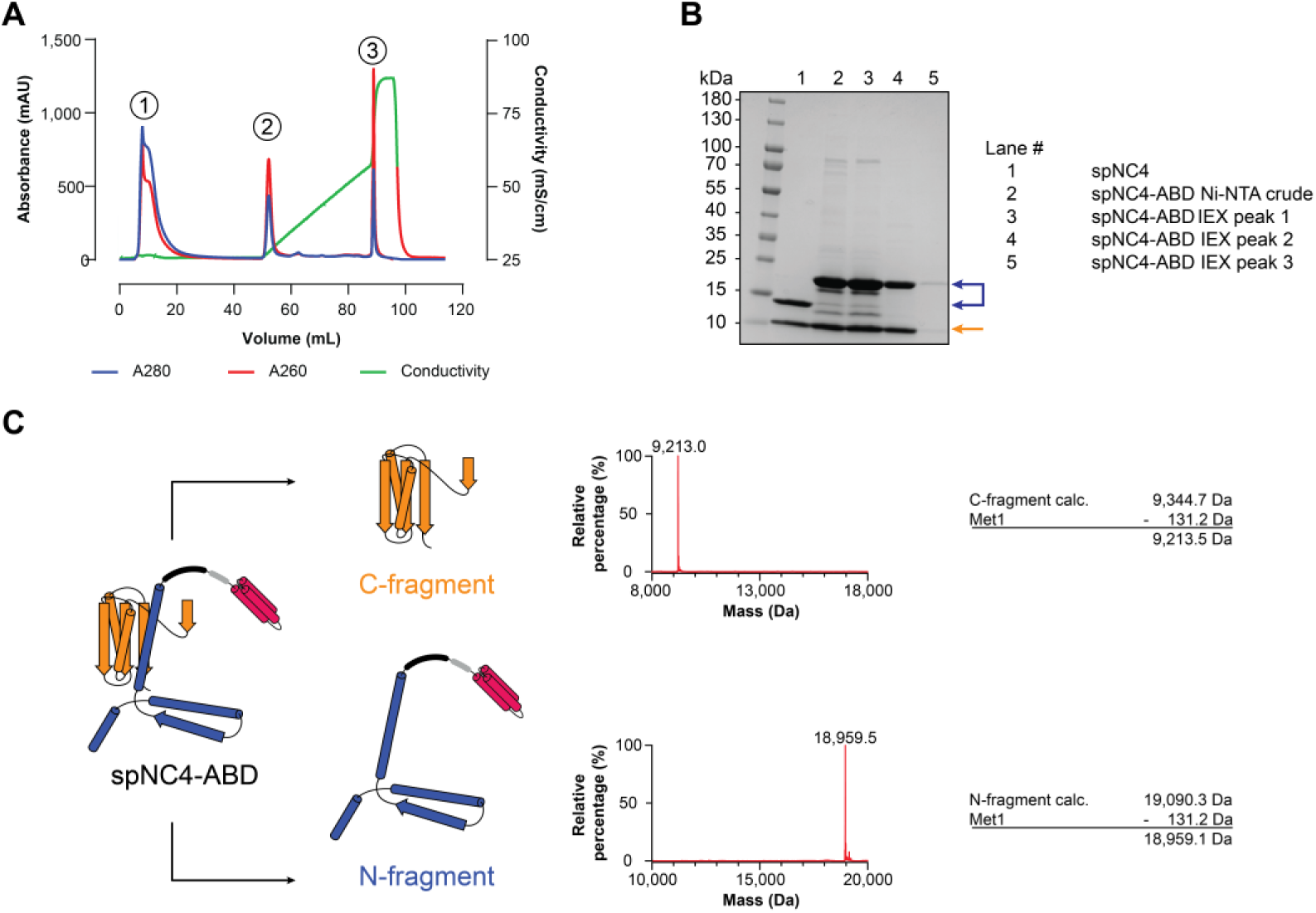
Characterization of the spNC-4-ABD variant. (**A**) Ion-exchange (IEX) chromatography of spNC-4-ABD showing the trace of the crude extract after metal-ion affinity chromatography. Absorbance was recorded at both 260 nm (red) and 280 nm (blue); conductivity is indicated by the solid green line. (**B**) Coomassie-stained SDS-PAGE of spNC-4 and spNC-4-ABD. The N and C fragments of spNC-4 and spNC-4-ABD are marked by blue and orange arrows, respectively. Lanes 3 to 5 correspond to peaks 1 to 3 from the IEX chromatogram (panel A). (**C**) Mass spectrometry analysis of the C (top) and N (bottom) fragments of spNC-4-ABD.

### MATERIALS AND METHODS

#### Molecular cloning

All PCR products were purified by DNA Clean & Concentrator-5 (D4003, Zymo Research) or Zymoclean Gel DNA Recovery Kit (D4001, Zymo Research). All plasmids were purified using ZR Plasmid Miniprep-Classic (D4015, Zymo Research). Plasmid sequences were confirmed by Sanger sequencing (Microsynth AG). All DNA oligos were purchased from Microsynth AG and are listed in Table S1.

<u>pMG-dB-spNC-4</u>: The DNA fragment encoding the second ribosomal binding site was prepared by assembly PCR using primers NT1, NT2, NT3, and NT4; the fragment encoding the C-terminal segment of NC-4 was amplified from pMG-dB-NC-4^1^ with primers NT5 and NT6. The resulting DNA fragments were assembled with primers NT3 and NT6. The full-length gene was digested with SacI and XhoI and ligated into a pMG-dB-NC-4 plasmid that had been digested with the same restriction enzymes to give plasmid pMG-dB-spNC-4.

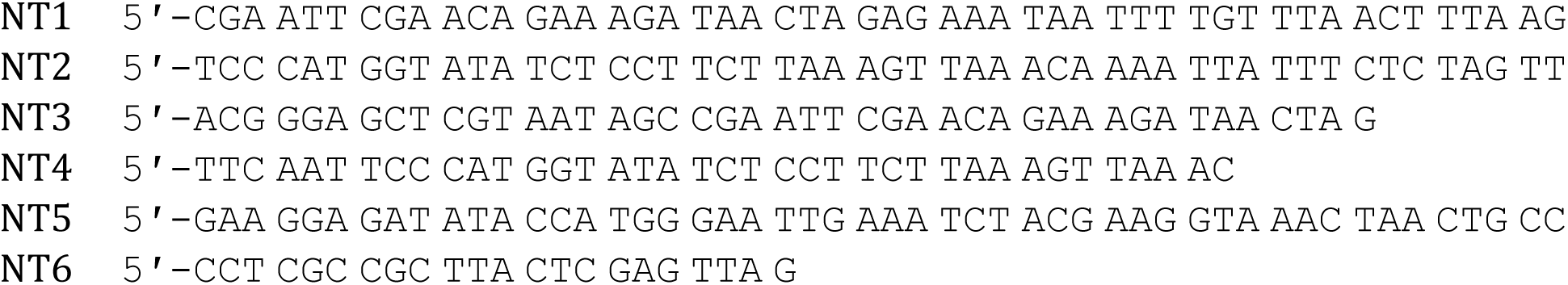

#### pMG-dB-spNC-4-GI and pMG-dB-spNC-4-ABD

GI construct: A synthetic DNA fragment encoding a glycosylation tag (11 aa), a spacer arm (7 aa), and a hexahistine tag fused to the C-terminus of the N-fragment of spNC-4 was obtained from Twist Biosciences.

Sequence of the GI-encoding fragment:

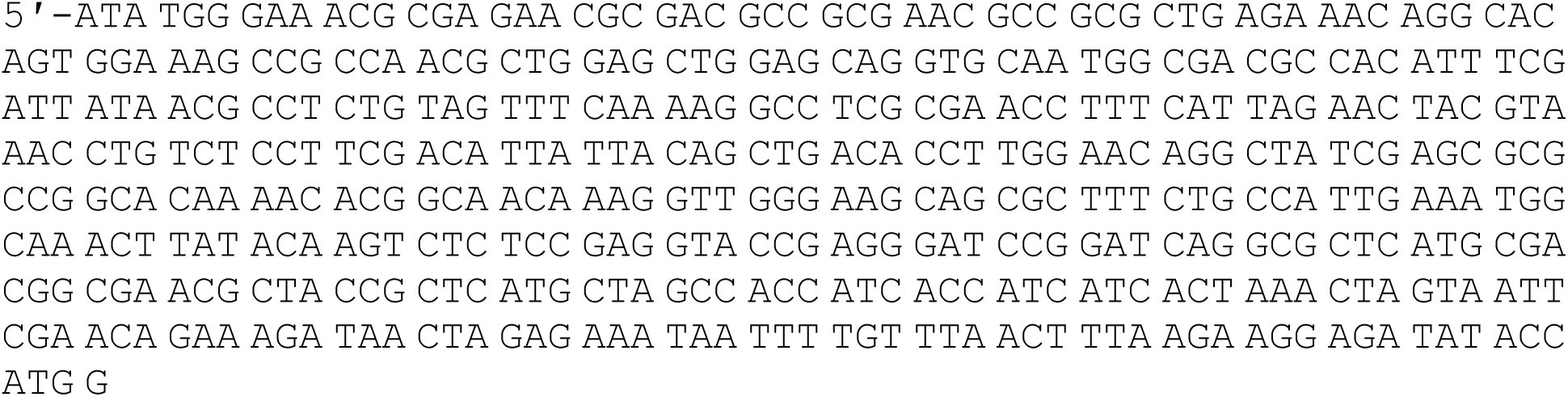

ABD construct: A synthetic DNA fragment encoding an antibody binding domain (ABD, 58 aa), a spacer (3 aa), and a hexahistine tag, also fused to the C-terminus of the N-fragment spNC-4, was obtained from Twist Biosciences.

Sequence of the ABD-encoding fragment:

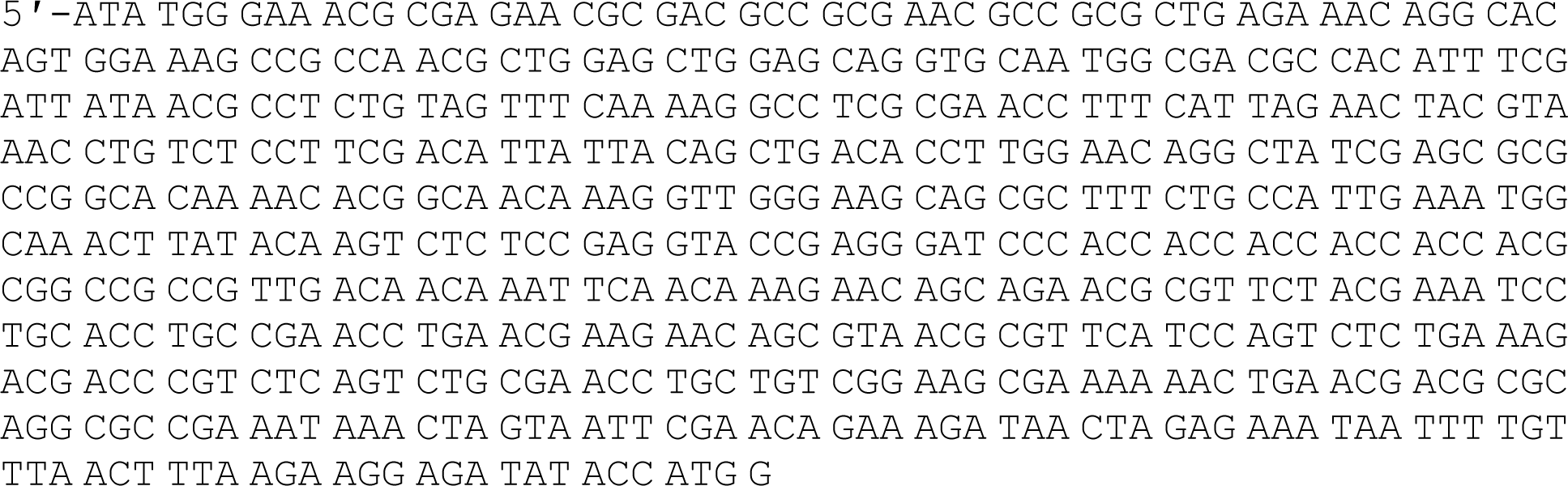

The 386 bp GI and 524 bp ABD inserts generated by digestion with XbaI and EcoRI at 37 °C for 3 hours were ligated into the correspondingly digested pMG-dB-spNC-4 backbone using T4 DNA ligase (NEB), following the manufacturer’s recommended protocol, to give the plasmids pMG-dB-spNC-4-GI and pMG-dB-spNC-4-ABD, respectively.

#### Production of NC-4 and spNC-4

Both NCs were produced in *E. coli* BL21(DE3)-gold cells. Briefly, 800 mL LB medium in 2-L Erlenmeyer flasks was inoculated with 8 mL overnight culture and incubated at 37 °C and 200 rpm until the OD600 reached 0.5–0.7. Protein production was induced by adding IPTG to a final concentration of 0.5 mM. Cultures were then grown for 18 hours at 25 °C for (NC-4) or at 30 °C (spNC-4) for 18 hours before harvesting by centrifugation at 6,000 x g and 15 °C for 20 min. The cell pellet from each 800-mL culture was resuspended in 20 mL LB medium, transferred, and divided into two 50-mL Falcon tubes. The medium used for transfer was removed by centrifugation at 4,000 x g and 15 °C for 10 min, decanted, and aliquots of the cell pellet were frozen in liquid nitrogen and stored at −20 °C until purification.

For purification, pellets corresponding to 400 mL culture were resuspended in 20 mL lysis buffer (50 mM sodium phosphate, pH 7.4, 1 M NaCl, 20 mM imidazole). The lysis buffer was supplemented with lysozyme (1 mg/mL). The mixture was incubated at room temperature with orbital shaking for 20 min. Cells were lysed by sonication (5 cycles of 1 min on, 1 min off, with amplitude = 80 and cycle = 60, UP200S sonicator, Hielscher Ultrasonics GmbH). Lysates were clarified by centrifugation at 8,500 x g and 15 °C for 25 min, and the supernatant was applied to 3 mL Ni(II)-NTA agarose resin in a gravity flow column. After 10 min incubation and washing with lysis buffer, the NCs were eluted with buffer (50 mM sodium phosphate buffer at pH 7.4) containing either 300 (spNC-4) or 500 mM (NC-4) imidazole. The eluted fractions were dialyzed overnight against storage buffer (50 mM sodium phosphate, pH 7.4, 200 mM NaCl, 5 mM EDTA) using 10 kDa MWCO SnakeSkin dialysis tubing (22 mm, Thermo Scientific). Protein capsids were purified by size exclusion chromatography at room temperature on a HiPrep 16/60 Sephacryl S-400 column (GE Healthcare Life Sciences) equilibrated in storage buffer, followed by anion exchange chromatography on a MonoQ 10/100 column (Pharmacia Biotech), employing a NaCl gradient from 200 to 1000 mM in storage buffer. Purified fractions were pooled, concentrated, aliquoted, and either analyzed immediately or stored at –80 °C after flash freezing in liquid nitrogen. Protein and RNA concentrations were determined by UV absorbance and deconvoluted using a previously reported method.^1^ Protein extinction coefficients were calculated with the ExPASy ProtParam tool.^4^

#### Negative-stain transmission electron microscopy (TEM)

Negative-stain TEM was performed as previously described.^1^ Briefly, TEM grids (#01814 F, Ted Pella, Inc.) were glow-discharged at 25 mA for 45 s using a Pelco easiGlow Glow Discharge Cleaning System. Following FPLC purification, grids were incubated with capsid solution (10 μM monomer in storage buffer containing 200 mM NaCl) for 1 min. The grids were then washed twice with doubly distilled water (ddH2O), and once with staining solution (2% wt/vol aqueous uranyl acetate, pH 4), followed by a 10 s incubation with staining solution. After drying, grids were imaged using a TFS Morgagni 268 microscope.

#### In vitro transcription of reference mRNAs

Reference messenger RNAs (mRNAs) samples were prepared as described previously.^1^ Briefly, DNA templates were generated by PCR amplification from plasmids pMG-dB-NC-4 and pMG-dB-spNC-4 using primers 1 (5′-GCG AAA TTA ATA CGA CTC ACT ATA G) and 2 (5′-CAA AAA ACC CCT CAA GAC CC) with the LongAmp Taq assay kit (#M0287, NEB). PCR-amplified products were gel-purified using the DNA Clean & Concentrator-5 kit. In vitro transcription was performed using T7 RNA polymerase (#EP0111, Thermo Scientific) following the manufacturer’s instructions. After transcription, template DNA was removed by RQ1 DNase digestion, and RNA was precipitated with isopropanol. RNA samples were purified twice by denaturing polyacrylamide electrophoresis (PAGE), using preparative gels (20 cm x 16 cm x 0.1 mm) composed of 5% polyacrylamide prepared in Tris/borate/EDTA (TBE) buffer containing 8 M urea. Polymerization was initiated with TEMED (8 μL per 10 mL gel solution) and 10% aq. ammonium persulfate (APS) (90 μL/10 mL gel solution). RNA bands were visualized by UV shadowing and excised with a scalpel. Gel pieces were crushed with a pipet tip, and the RNA was eluted overnight at room temperature in water containing 0.3 M NaCl. The next day, the RNA was purified by ethanol precipitation and dissolved in water. RNA quality and purity were assessed by measuring A_260_/A_280_ and A_260_/A_230_ absorbance ratios (for pure RNA, both ratios are ≥2.0) and by analytical PAGE. RNA concentrations were determined using the Qubit RNA HS assay (#Q32852, Invitrogen).

#### Extraction of NC RNA and RT-qPCR

RNA extraction and RT-qPCR were performed as previously described.^3^ Briefly, RNA was extracted from 100-or 200-µL aliquots of purified NCs containing 5–10 µg total RNA using the RNeasy Mini kit (#74104, QIAGEN) following the manufacturer’s instructions. RNA standards were prepared by in vitro transcription as described above. After confirming RNA purity by absorbance measurements, concentrations of extracted RNAs and in vitro-transcribed standards were determined using the Qubit RNA HS Assay. For capsid cDNA synthesis, reverse transcription was performed with primer 3 (5′-CCA AGG GGT TAT GCT AGT TAT TGC TCA GC) and SuperScript III reverse transcriptase (#18080044, Invitrogen) according to the manufacturer’s protocol. After the reverse transcription reaction, RNase H (#18021014, Invitrogen) was added to degrade RNA transcripts. Immediately afterward, dilutions of the cDNA were mixed with KOD SYBR qPCR Master Mix (#QKD-201, TOYOBO), primers 4 (5′-TGT GAG CGG ATA ACA ATT CCC CTC) and 5 (5′-GGG TTA TGC TAG TTA TTG CTC AGC G), and ROX reference dye per the manufacturer’s guidelines. qPCR was carried out over 40 PCR cycles on a StepOnePlus thermocycler (Applied Biosystems) employing the thermocycler-specific conditions provided in the qPCR mix manual. Absolute amounts of full-length cDNA were determined using standard curves prepared with cDNA originating from highly pure in vitro-transcribed reference RNAs.

#### Analytical urea PAGE

Analytical urea polyacrylamide gel electrophoresis (PAGE) was performed as described previously.^3^ Analytical gels (8.3 cm x 7.3 cm x 0.1 mm) were prepared in TBE buffer containing 8 M urea and 8% polyacrylamide. Polymerization was initiated with TEMED (8 μL/10 mL gel solution) and APS (10% in water, 90 μL per 10 mL gel solution). Gels were loaded with equal volumes of extracted RNA. The NC genome was selectively visualized using the fluorogenic dye DFHBI-1T (#446461, United States Biological), which fluoresces upon binding to the Broccoli aptamer in the BoxBr tags.^3^ Total RNA was stained separately with GelRed (#41002, Biotium).

#### CryoEM

For cryoEM, spNC-4 was adjusted to a final concentration of approximately 2 mg/mL. Copper-supported holey carbon grids (R2/2, Cu 400, Quantifoil) were treated by negative glow discharge at 15 mA for 15 s using a Pelco easiGlow Glow Discharge Cleaning System. 3.5 µL of the sample were applied to the grids and blotted for 12 s at a blot force of 25, 100% humidity, and 22 °C using a Vitrobot (FEI). Grids were plunged in liquid ethane and stored in liquid nitrogen.

Data were acquired on a Titan Krios microscope fitted with a Falcon III detector (FEI), with 40-frame movies collected at a total dose of 60 e⁻/Å² and a magnification of 130,000× (1.1 Å pixel size). Data were recorded in electron counting mode, with defocus values between –0.8 and –2.7 µm.

All single-particle reconstructions were processed using Relion 3.0.^5^ Motion correction was performed with MotionCor2^6^ and contrast transfer function (CTF) parameters were estimated with GCTF.^7^ Well-defined 2D classes were used to generate 3D models de novo under icosahedral symmetry constraints. The best 3D classes were further refined with masks applied to the capsid shells, masking out the interior.

Model building and refinement were performed using Coot (v0.8.9.2),^8^ Phenix (v1.18),^9^ and PyMOL (v2.0). Density maps derived from Relion postprocessing and Phenix autosharpening were used for model construction. The model was built based on the published NC-4 structure (PDB: 7a4j). Initial models were built in the asymmetric unit and refined before applying symmetry expansion, after which the complete capsids were refined under non-crystallographic symmetry constraints. Detailed data collection and model statistics are summarized in Table S1.

#### Sample preparation for solid-state NMR

The spNC-4 and NC-4 nucleocapsids were both produced in *E. coli* BL21(DE3)-gold cells. Three-liter Erlenmeyer flasks containing 1 L LB medium were inoculated with 10 mL overnight cultures and incubated at 37 °C and 200 rpm until OD600 reached approximately 0.5. Cells were pelleted by centrifugation at 5,000 x g and 15 °C for 30 min, washed with nitrogen-and carbon-free M9 salt solution, and pelleted again by centrifugation at 5,000 x g and 15 °C for 15 min. Cells were then resuspended in M9 media supplemented with ^13^C-labeled glucose and ^15^N-labeled ammonium chloride and incubated at 37 °C to allow growth recovery and clearance of unlabeled metabolites, as previously described.^10^ Protein production was induced by adding IPTG to a final concentration of 0.5 mM. Cells producing NC-4 were cultured at 20 °C, while those expressing spNC-4 were cultured at 30 °C for 18 hours. Cells were harvested by centrifugation at 6,000 x g and 15 °C for 20 min. The pellet from one 1-L culture was resuspended in 20 mL LB medium, split into two 50-mL Falcon tubes. The medium used for transfer was removed by centrifugation at 4,000 x g and 15 °C for 10 min, decanted, and aliquots of the cell pellet were flash-frozen in liquid nitrogen and stored at -20 °C until purification.

For purification, pellets corresponding to 500 mL culture volume were resuspended in 20 mL lysis buffer (50 mM sodium phosphate, pH 7.4, 1 M NaCl, 20 mM imidazole) supplemented with lysozyme (1 mg/mL) and incubated at room temperature with orbital shaking for 20 min. Cells were lysed by sonication (5 cycles of 1 min on, 1 min off, with amplitude = 80 and cycle = 60, UP200S sonicator, Hielscher Ultrasonics GmbH). Lysates were clarified by centrifugation at 7,000 x g and 15 °C for 25 min, and the supernatant was applied to 3 mL of Ni(II)-NTA agarose resin in a gravity flow column. After 10 min incubation and washing with lysis buffer, the nucleocapsids were eluted with buffer containing 50 mM sodium phosphate, pH 7.4, 300 mM NaCl, and 500 mM imidazole. Eluted fractions were concentrated and buffer-exchanged into storage buffer (50 mM sodium phosphate, pH 7.4, 200 mM NaCl, 5 mM EDTA) using Amicon Ultra-15 centrifugal filter units (100 kDa MWCO, Merck Millipore). Protein capsids were further purified by size exclusion chromatography at room temperature using a HiPrep 16/60 Sephacryl S-400 column (GE Healthcare Life Sciences) equilibrated in storage buffer, followed by anion exchange chromatography on a MonoQ 10/100 column (Pharmacia Biotech) at room temperature using a NaCl gradient from 200 to 1,000 mM in storage buffer. Purified fractions were pooled and concentrated using Amicon Ultra-15 units (100 kDa MWCO). Protein and RNA concentrations were determined by UV absorbance and deconvoluted using a previously reported protocol.^3^ Extinction coefficients for proteins were calculated using the ExPASy ProtParam tool.^4^

Prior to NMR sample preparation, protein samples were concentrated using Amicon centrifugal filters and supplemented with sodium trimethylsilylpropanesulfonate (DSS) dissolved in water for chemical shift referencing and sodium azide (NaN₃) to prevent microbial growth. Samples were packed into 3.2 mm or 0.7 mm rotors using NMR filling tools^12^ by ultracentrifugation overnight at 200,000 x g and 4 °C.

#### Solid-state NMR spectroscopy

Solid-state NMR spectra were recorded on a Bruker Avance III wide-bore 850 MHz spectrometer. Protein samples were spun 3.2 mm rotors at 17 kHz MAS for carbon-detected experiments and in 0.7 mm rotors at 100 kHz for proton-detected experiments. The sample temperature was maintained at 3-5 °C during carbon-detected experiments and 19-22 °C during proton detection. Spectral processing included referencing to DSS, zero filling to double the number of data points, and application of a shifted sine-bell apodization function with Bruker TOPSPIN software version 3.5. Spectra were analyzed with CcpNmr version 2.^2^

Sequential assignment of spNC-4 was achieved by a ’backbone walk’ strategy based on reported methods,^13,14^ utilizing NCACB, NCACX, CANCO, and NCOCX spectra. Additional NcoCACB and CANcoCA experiments facilitated confidential sequential assignment progression. Two-dimensional ^13^C-^13^C DARR, NCA, and NCO spectra were used for orientation, monitoring assignment progress, and assessing spectral quality and sample stability throughout the measurement. Proton-detected three-dimensional spectra (hCANH, hNCAH, hNcoCAH, and hCAcoNH) were recorded as previously reported.^15^ Here, two-dimensional hNH and hCH experiments aided assignment and sample stability monitoring. Amide and HA protons were assigned by a sequential walk via CA and N resonances. Assignments from spNC-4 were transferred to NC-4, reducing the number of experiments needed for NC-4 assignment. A comprehensive list of recorded NMR spectra is provided in Table S2, whereas experimental parameters are summarized in Tables S2 and S3 for spNC-4 and NC-4, respectively. Assignment data have been deposited in the Biological Magnetic Resonance Data Bank under accession numbers XXXXXX (spNC-4) and YYYYYY (NC-4).

#### Production and purification of glucosylated spNC-4 constructs

For glycosylation of spNC-4 nucleocapsids, 2-L Erlenmeyer flasks were filled with 800 mL LB media (LB Broth, Miller– DIFCO #244610) containing 100 μg/mL sodium ampicillin and 25 μg/mL chloramphenicol. Flasks were inoculated with a single colony of *E. coli* BL21-Gold (DE3) strains co-transformed with pMA933 (a pACYC-DUET plasmid with the ApNGT gene under the control of a lacUV5 promoter and the chloramphenicol resistance gene CmR) along with the spNC-4 or spNC-4-GI plasmids. Cultures were grown at 37 °C with shaking at 230 rpm. When the OD600 reached 0.6– 1.0, protein expression was induced by adding 2 μL of a 1M IPTG solution to give a final concentration of 0.4 mM. After incubating for 18 hours at 25 °C and 230 rpm, cells were harvested by centrifugation at 5,000 x g and 4 °C for 45 min. Pellets were stored at −20 °C until purification as described above for spNC-4. Purified samples were subsequently analyzed by mass spectrometry.

#### Production and purification of spNC-4-ABD constructs

The spNC-4-ABD construct was expressed and purified using the same procedure as described for NC-4 and spNC-4, with one modification: following Ni(II)-NTA affinity chromatography, the eluate was directly subjected to anion exchange chromatography without intermediate size exclusion chromatography. All other steps, including expression conditions, lysis, affinity purification, and dialysis, were performed identically. For cellular uptake experiments, both spNC-4 and spNC-4-ABD were expressed and purified using this procedure. Both proteins were used immediately after purification for cellular uptake experiments, without freezing.

#### Cell culture

SKBR-3 cells were maintained in McCoy’s 5A medium (Sigma #M8403). MCF-7 cells were maintained in Iscove’s modified Dulbecco’s medium (Sigma #I3390). In all cases, media were supplemented with 10% fetal bovine serum (FBS), 2 mM L-glutamine, 2 mM GlutaMAX, and 1 μg/mL gentamicin. Cells were seeded at 1 x 10^6^ cells per T75 flask and incubated at 37 °C with 5% CO_2_ for 2–3 days.

#### Fluorescent labeling of spNC-4 cages

Labeling of spNC-4 and spNC-4-ABD cages with Atto565-NHS-ester was performed by mixing purified protein in storage buffer with a 20 mM dye solution in DMSO at a dye-to-cage molar ratio of 10:1, based on the assumption that each protein cage consists of 240 subunits. The mixture was incubated in the dark at room temperature for 18 hours. Unreacted dye was removed using a PD Minitrap G-10 desalting column equilibrated with PBS, and the labeled protein was collected for further analysis. Labeling efficiency and protein concentration were assessed by UV-Vis absorbance spectroscopy. The absorbance at 280 nm was corrected for the dye’s contribution using the manufacturer’s correction factor (CF_280_ = 0.12), and this corrected value was used to calculate protein concentration based on the molar extinction coefficients of 20,970 M⁻¹·cm⁻¹ for spNC-4 and 22,460 M⁻¹·cm⁻¹ for spNC-4-ABD. The dye concentration was derived from the absorbance at 564 nm, using the dye’s molar extinction coefficient of 120,000 M⁻¹·cm⁻¹. The degree of labeling (DOL), or the average number of dye molecules per cage, was calculated by dividing the dye concentration by the protein concentration adjusted for the 240-subunit cage assembly, resulting in typical DOL values of approximately 1.04 dye per cage for spNC-4 and 3.39 dyes per cage for spNC-4-ABD. The higher labeling efficiency observed for spNC-4-ABD can be attributed to six additional surface-exposed lysine residues, which are more accessible to the dye and thus facilitate greater conjugation.

#### Cellular uptake by flow cytometry

Cells were seeded at a density of 2 x 10^5^ cells per well in a 24-well plate containing 500 μL of culture medium and allowed to recover for 24 hours at 37 °C and 5% CO_2_. Atto565-labeled protein cages and Herceptin were sterilized separately by filtration through 0.22 μm membranes, and stock solutions were prepared in sterile storage buffer (50 mM sodium phosphate, pH 8, 200 mM NaCl, 5 mM EDTA). Prior to treatment, culture medium was removed and cells were washed once with 500 μL of Dulbecco’s Modified Eagle Medium (DMEM, Invitrogen) supplemented with 1 μg/mL gentamicin but without fetal bovine serum (FBS). In each well, 50 μL of labeled protein cage solution in PBS was added to 500 μL of serum-free DMEM, yielding a final volume of 550 μL. This dilution resulted in final dye concentrations ranging from 46.7 to 49.7 nM, with corresponding cage concentrations of 45.1 nM for spNC-4 (approximately 10.8 μM subunit concentration) and 14.7 nM for spNC-4-ABD (approximately 3.5 μM subunit concentration). Where indicated, Herceptin was added at a 10:1 molar ratio relative to the protein cage, resulting in final antibody concentrations of 436.4 nM for spNC-4 + Herceptin and 145.5 nM for spNC-4-ABD + Herceptin. The final concentration of each component in the different samples was as follows:

| Sample | Final dye (nM) | Final cage (nM) | Final Herceptin (nM) |
| --- | --- | --- | --- |
| spNC-4 | 46.7 | 45.1 | 0.0 |
| spNC-4-ABD | 49.7 | 14.7 | 0.0 |
| spNC-4 + Herceptin | 46.7 | 45.1 | 436.4 |
| spNC-4-ABD + Herceptin | 49.7 | 14.7 | 145.5 |

Cells were incubated for 2 hours in 5% CO_2_ at 37 °C before washing with 500 μL ice-cold PBS and trypsinization (0.05% trypsin-EDTA (ThermoFisher), 5 min at 37 °C). Cells were collected in cold culture medium (500 μL McCoy’s 5A or Iscove’s modified Dulbecco’s medium) and washed twice with 5 mL ice-cold PBS. All centrifugation steps were carried out at 500 x g for 5 min at room temperature. The cells were resuspended in flow cytometry buffer (500 μL PBS with 3% FBS). Samples were transferred to a 5-mL polystyrene round-bottom tube with a cell-strainer cap (Flacon #352235) and analyzed on an LSRFortessa flow cytometer (BD Biosciences). Each condition was tested in two replicate wells containing identical protein samples.

### PROTEIN SEQUENCES

NC-4

MGNARTRRRERRAEKQAQWKAANAGAGAGAMATPHFDYNASVVSKGLANLSLELRKPVSFDIIT ADTLEQAIERAGTKHGNKGWEAALSAIEMANLYKSLRGTEHHHHLHGSSIEIYEGKLTAEGLRF GIVASRFNHTLVDRLVEGAIDCIVRHGGRGEDITLVRVPGAWEIPVAADELARKEDIDAVIAFG DLIRG

spNC-4 N-fragment

MGNARTRRRERRAEKQAQWKAANAGAGAGAMATPHFDYNASVVSKGLANLSLELRKPVSFDIITADTLEQAIERAGTKHGNKGWEAALSAIEMANLYKSLRGTEHHHHLHGSS

spNC-4 C-fragment

MGIEIYEGKLTAEGLRFGIVASRFNHTLVDRLVEGAIDCIVRHGGRGEDITLVRVPGAWEIPVAADELARKEDIDAVIAFGDLIRG

spNC-4-GI N-fragment

MGNARTRRRERRAEKQAQWKAANAGAGAGAMATPHFDYNASVVSKGLANLSLELRKPVSFDIITADTLEQAIERAGTKHGNKGWEAALSAIEMANLYKSLRGTEGSGSGAHATANATAHASHHHHHH

spNC-4-GI C-fragment

MGIEIYEGKLTAEGLRFGIVASRFNHTLVDRLVEGAIDCIVRHGGRGEDITLVRVPGAWEIPVAADELARKEDIDAVIAFGDLIRG

spNC-4-ABD N-fragment

MGNARTRRRERRAEKQAQWKAANAGAGAGAMATPHFDYNASVVSKGLANLSLELRKPVSFDIITADTLEQAIERAGTKHGNKGWEAALSAIEMANLYKSLRGTEGSHHHHHHAAAVDNKFNKEQQNAFYEILHLPNLNEEQRNAFIQSLKDDPSQSANLLSEAKKLNDAQAPK

spNC-4-ABD C-fragment

MGIEIYEGKLTAEGLRFGIVASRFNHTLVDRLVEGAIDCIVRHGGRGEDITLVRVPGAWEIPVAADELARKEDIDAVIAFGDLIRG

## Notes

### Competing Interest Statement

The authors have declared no competing interest.

